# Pericytes control vascular stability and auditory spiral ganglion neuron survival

**DOI:** 10.1101/2022.09.27.509698

**Authors:** Yunpei Zhang, Lingling Neng, Kushal Sharma, Zhiqiang Hou, Anatasiya Johnson, Junha Song, Alain Dabdoub, Xiaorui Shi

## Abstract

The inner ear has a rich population of pericytes, a multi-functional mural cell essential for sensory hair cell heath and normal hearing. However, the mechanics of how pericytes contribute to the homeostasis of the auditory vascular-neuronal complex in the spiral ganglion is not yet known. In this study, using an inducible and conditional pericyte depletion mouse (*Pdgfrb*^*CreERT2+/-*^; *ROSA26*^*iDTR+/-*^) model, we demonstrate, for the first time, that pericyte depletion causes loss of vascular volume and spiral ganglion neurons (SGNs) and adversely affects hearing sensitivity. Using an *in vitro* trans-well co-culture system, we show pericytes markedly promote neurite and vascular branch growth in neonatal SGN explants and adult SGNs. The pericyte-controlled neural growth is strongly mediated by pericyte-released exosomes containing vascular endothelial growth factor-A (VEGF-A). Treatment of neonatal SGN explants or adult SGNs with pericyte-derived exosomes significantly enhances angiogenesis, SGN survival, and neurite growth, all of which were inhibited by a selective blocker of the VEGF receptor 2 (Flk1). Our study demonstrates that pericytes in the adult ear are critical for vascular stability and SGN health. Cross-talk between pericytes and SGNs via exosomes is essential for neuronal and vascular health and normal hearing.

## Introduction

The inner ear is dense in vascular beds. Microvascular networks are situated in different locations of the cochlea (Shi, 2011). The major microvascular network is located in the cochlear lateral wall, receiving ∼ 80% of cochlear blood flow (Gyo, 2013). Blood flow to the cochlear lateral wall is essential for cochlear homeostasis and is particularly important for generating the endocochlear potential necessary for sensory hair cells (HCs) transduction (Hibino et al., 2010). The next largest microvascular network is situated around the region of the spiral ganglia neurons (SGNs), comprising ∼ 19% −24% of blood flow and is critical for neural activity (as illustrated in **Fig. 1**) (Angelborg et al., 1984; Gyo, 2013; Nakashima et al., 2001). In particular, the vascular network of the spiral ganglion directly delivers nutrients and growth hormones to SGNs. Defects in vascular structure and function would significantly and quickly affect the viability of vulnerable neural tissue. Loss of SGNs is commonly associated with different types of hearing loss (Leake et al., 2020; Viana et al., 2015).

**Figure 1.**
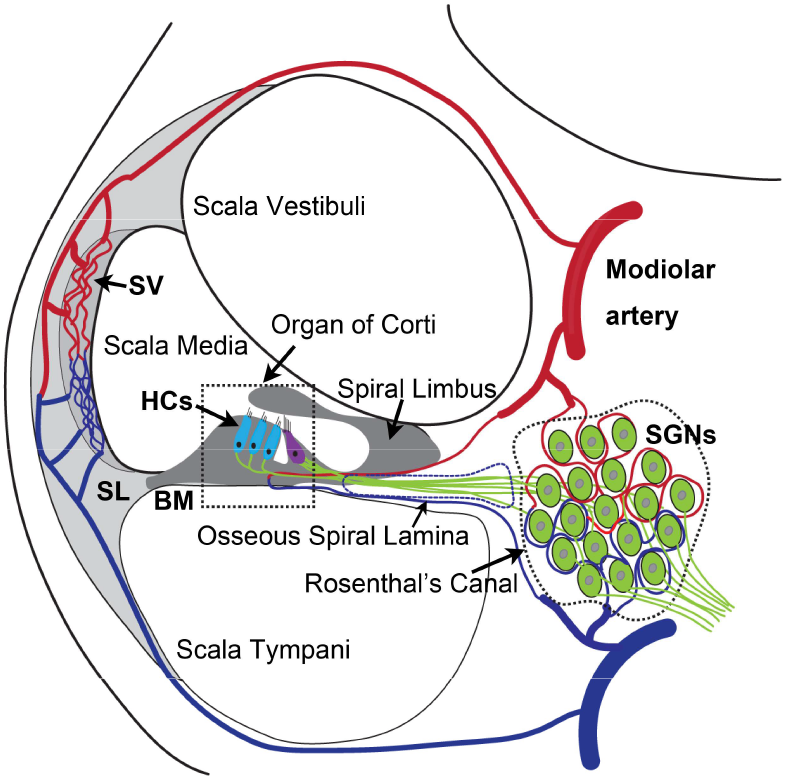
The anatomy and blood supply of cochlea. Two major microvascular networks in the cochlea include the network in the cochlear lateral wall and network in the region of SGNs - Blood vessels penetrate the SGNs and directly supply nutrients to the neurons. SV, stria vascularis; SL, spiral ligament; HCs, hair cells; BM, basilar membrane; SGNs, spiral ganglion neurons.

Pericytes are a type of mural cell that encircle endothelial cells of capillaries, pre-capillary arterioles, and post-capillary venules (Armulik et al., 2011). They extend their processes along and around these vessels and play critical roles in regulating vascular morphogenesis and function (Attwell et al., 2016; Birbrair, 2018). Normal function of pericytes and adequate blood supply are crucial for providing vital oxygen, ions, and glucose to meet organ metabolic needs (Shaw et al., 2018). Pericyte degeneration or loss has been identified as a major pathology in many diseases, including ischemic stroke (Greif & Eichmann, 2014) and brain trauma (Zehendner et al., 2015), myocardial Infarction (O’Farrell & Attwell, 2014), diabetic retinopathy (Pfister et al., 2008), and neurodegeneration diseases such as Alzheimer’s disease, Parkinson’s disease, and Huntington’s disease (Sweeney et al., 2018). Pericytes are known to display phenotypic and functional heterogeneity, which is highly associated with organ function (Dias Moura Prazeres et al., 2017). While little is known about the role of pericytes in the inner ear; nothing is known about how pericyte loss affects peripheral neuron health.

SGN cell bodies, located in Rosenthal’s canal, extend distal processes radially outward into the spiral lamina toward sensory hair cells in the organ of Corti, and central processes project into the auditory nerve (as illustrated in **Fig. 1**) (Nayagam et al., 2011). The microvascular network, located in the spiral ganglion region, forms radial vascular twigs that directly supply nutrients to the SGNs. Although the volume of blood flow to the spiral limbus and SGN is lower than to the cochlear lateral wall, it is critical for neuronal activities. Constant sound stimulation to the inner ear imposes a high energy demand on neurons, requiring rapid delivery of oxygen and glucose. Previously we demonstrated the microvascular network in the spiral ganglion region is richly populated by pericytes (Jiang et al., 2019). Pericytes actively communicate with SGNs as evidenced by the SGNs taking up pericyte-released particles (Jiang et al., 2019). However, the role of pericytes in the SGN viability and stability of the vascular beds is not yet known. In this study, using an inducible and conditional genetic pericyte ablation model in combination with two newly established co-culture models, we demonstrate that pericyte loss *in vivo* causes loss of vessel volume and SGNs in adult mice. *In vitro*, we demonstrate vigorous vascular and neuronal growth in the neonatal SGN explants in the presence of exogenous pericytes. Increased SGN survival and neurite growth were also observed in adult SGNs. Most interestingly, we find the promotion of vascular and neuronal growth is mediated by pericyte-released exosomes, nano-sized 50 to 150 nm extracellular vesicles (EVs), containing vascular endothelial growth factor-A (VEGF-A). VEGF-A binding to VEGFR2 (Flk1) expressed in recipient cells, including endothelial cells and SGNs, controlled vascular and neuronal growth. Our data demonstrate for the first time that loss of pericytes leads to a reduction in vascular density and degeneration of SGNs, underscoring the vital role of pericytes in vascular and auditory peripheral neural health.

## Results

### Loss of pericytes leads to reduced capillary volume in the spiral ganglion region *in vivo*

To investigate whether pericyte loss affects vascular stability in the spiral ganglion in adult mice, we created an inducible pericyte depletion mouse model (*Pdgfrb*^*CreERT+/-*^; *R26*^*iDTR+/-*^) by crossing *Pdgfrb*^*CreERT2+/-*^ transgenic mice with *ROSA26*^*iDTR+/+*^ mice, an inducible daiphtheria toxin receptor (iDTR) mouse line carrying Cre-dependent simian DTR, which leads to cell death with the administration of the diphtheria toxin (DT) (Buch et al., 2005; Zhang et al., 2021) (as illustrated in **Fig. 2A**). This *Pdgfrb*^*CreERT+/-*^; *R26*^*iDTR+/-*^ mouse line at one month of age received tamoxifen (TAM) for three days to induce the expression of Cre recombinase. For ablation of pericyte, DT was administrated by daily intraperitoneal injection at 10 µg/kg for 4 consecutive days beginning 1 day after TAM (Zhang et al., 2021) (as illustrated in **Fig. 2B**). The *Pdgfrb*^*CreERT2-/-*^; *R26*^*iDTR+/-*^ mice from same litters of *Pdgfrb*^*CreERT2+/-*^; *R26*^*iDTR+/-*^ mice were treated with TAM and DT and constituted a control group. The cellular location of Cre recombinase expression under the *pdgfrb*-promotor was confirmed by crossing the *Pdgfrb*^*CreERT2*^ mice with *ROSA26*^*tdTomato*^ mice. As expected, the tdTomato fluorescence signal (Greenhalgh et al., 2013) co-localized exclusively with the immunofluorescence signal for the pericyte marker protein, PDGFRβ (green) in the vascular networks, as shown in **Fig. 2C and D**.

**Figure 2.**
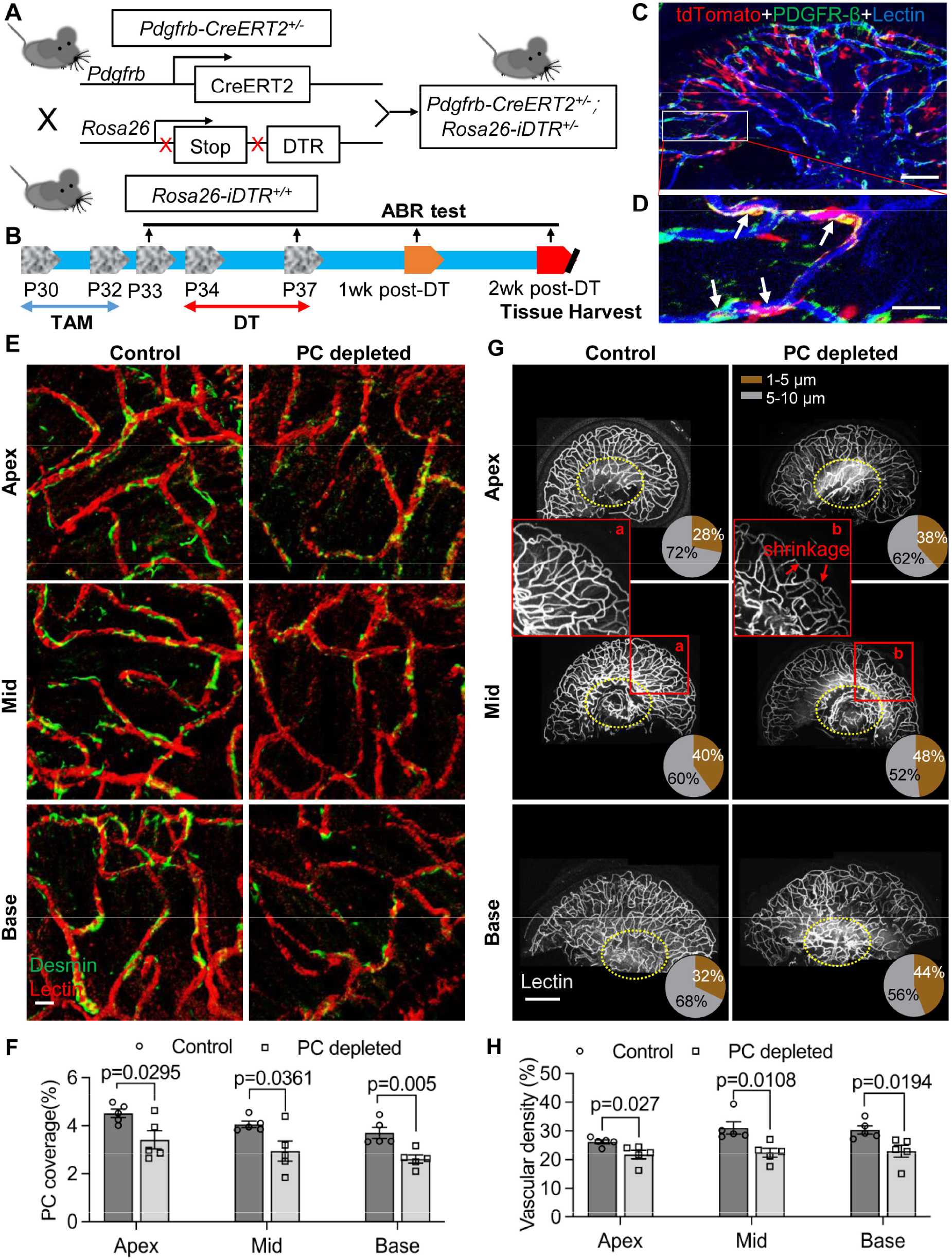
Pericyte-depletion induces vascular regression in the region of SGN peripheral nerve fibers in adult mice. (A) Schematic of a pericyte-depletion mouse model incorporating an inducible Cre-loxP system. (B) The diagram shows the timeline of tamoxifen and DT administration and the time point of ABR test and tissue harvest. (C) Co-localization of PDGFRβ-Cre (tdTomato) and immune-labeled PDGFRβ (green) signals in the pericytes of *Pdgfrb*^*CreERT2*^; *ROSA26*^*tdTomato*^ mice. (D) A high magnification image further showing co-localization of the Cre and PDGFRβ fluorescence signals (arrows). (E) Representative figures show pericyte coverage in the region of the spiral limbus in DT-treated control iDTR (left) and DT-treated *Pdgfrb*^*CreERT2*^; *ROSA26*^*iDTR*^ mice (right) 2 weeks after DT injection. DT injection significantly leads to loss of pericyte coverage. (F) Pericyte density was significantly reduced in the *Pdgfrb*^*CreERT2*^; *ROSA26*^*iDTR*^ mice at the apical, middle, and basal turn relative to density in the control of iDTR mice (n=5, P_Apex_=0.0295, P_Mid_=0.0361, P_Base_=0.005, unpaired t test). (G) Representative figures show the capillaries of the spiral lamina in control iDTR (left) and *Pdgfrb*^*CreERT2*^; *ROSA26*^*iDTR*^ (right) mice, with the distribution of vessel diameter shown in πcharts, and the location of SGNs shown in ellipses. (H) Total vascular density in the spiral limbus and lamina is significantly reduced in the pericyte-depleted mice 2 weeks after DT injection (n=5, P_Apex_=0.027, P_Mid_ =0.0108, P_Base_=0.0194, unpaired t test). Loss of vascular volume with pericyte depletion is better seen in the high magnification image inserts in panel F (a), (b). Data are presented as the mean ± SEM. Scale bars: C, 50 µm; D, 10 µm; E, 20 µm; F, 200 µm.

Our results showed, relative to control mice, pericyte distribution (labeled with desmin) in the vascular networks (labeled with Alexa Fluor™ 649 conjugate lectin, the most widely used fluorescence dye to visualize blood vessels (Meyer et al., 2008) of the spiral ganglion region was markedly reduced at all cochlear turns two weeks after DT treatment in the CreERT2/iDTR mice (n=5, P_Apex_=0.0295, P_Mid=_0.0361, P_Base_=0.005, unpaired t-test) (**Fig. 2E and F**). In addition, the vascular volume in the spiral ganglion region was also notably reduced. **Fig. 2G** has representative confocal images showing the pattern of blood vessel distribution in control and pericyte-depleted groups. **Fig. 2H** shows total vascular density in the spiral lamina is significantly reduced in the pericyte-depleted mice two weeks after DT injection (n=5, P_Apex_=0.027, P_Mid_=0.0108, P_Base_=0.0194, unpaired t-test). In addition, vascular shrinkage, a sign of degeneration, was frequently observed (as highlighted under high magnification, **Figure 2G**). Our results clearly indicate that pericytes are essential for vascular stability.

### Pericytes are critical for hearing sensitivity, as well as SGN function and survival *in vivo*

We further assessed hearing function with the auditory brainstem response (ABR) in pericyte-depleted mice. As shown in **Fig. 3A**, we found the control mice showed no significant hearing threshold change after DT injection. In contrast, the hearing threshold in pericyte-depleted animals was significantly elevated at 1 week after DT injection and persisted to two weeks after DT injection (n=10, P<0.0001, two-way ANOVA), consistent with our previous observation (Zhang et al., 2021).

**Figure 3.**
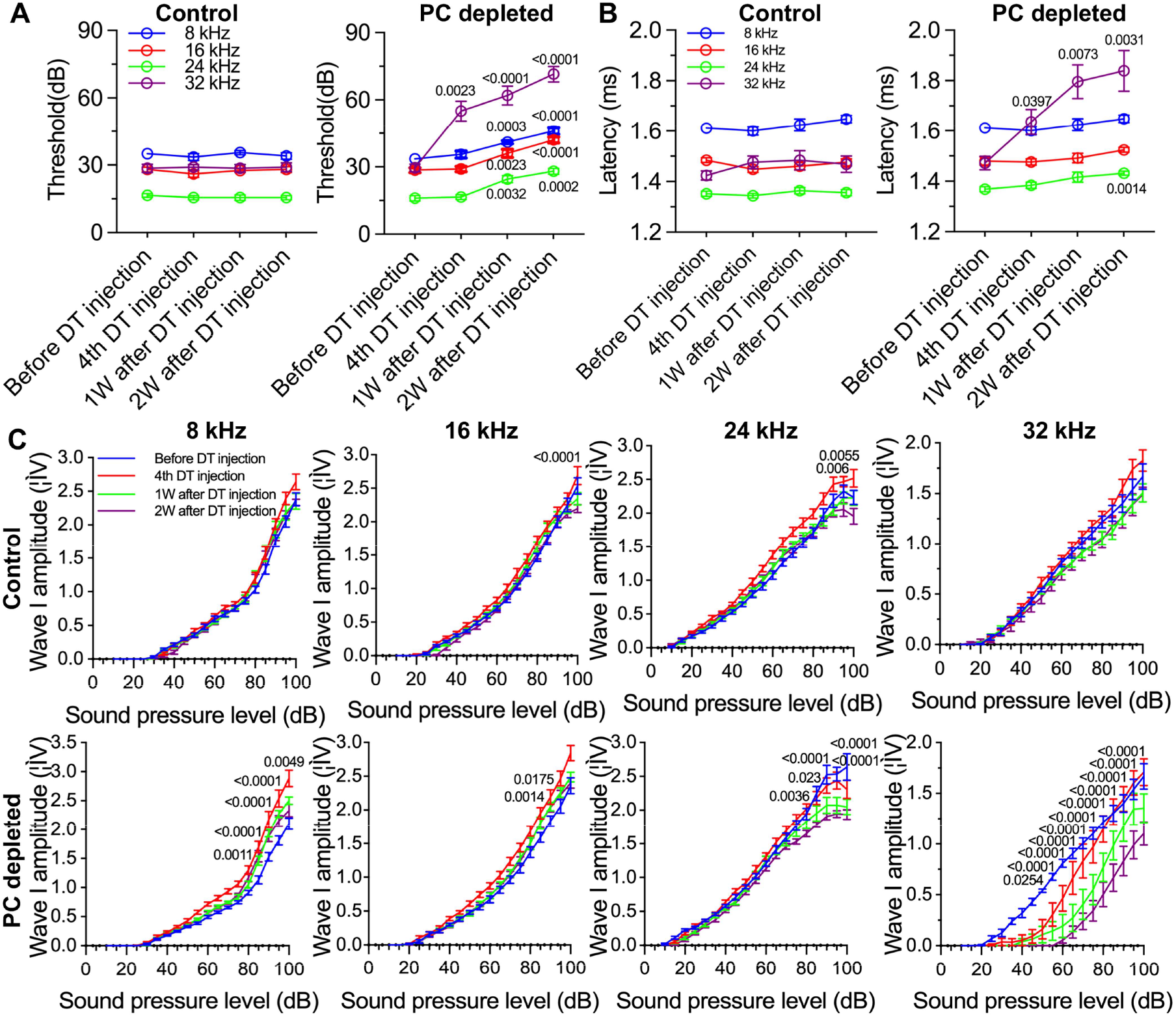
The depletion of pericytes led to hearing loss, increased latency and reduced amplitude of wave I. (A) The control mice showed no significant hearing threshold change after DT injection (n=10, P=0.9651). In contrast, the hearing threshold in pericyte-depleted animals was significantly elevated at 1 week after DT injection and persisted to two weeks after injection (n=10, P<0.0001). Two-way ANOVA, followed by Dunnett’s multiple comparison test, individual P values of different time points versus before DT injection are labeled on the graph. (B) The control mice showed no significant wave I latency change after DT injection (n=10, P=0.3576). In pericyte-depleted animals, the latency was significantly delayed at high frequency (32 kHz) started from 4^th^ DT injection, and low frequency (8 kHz) two weeks after DT injection (n=10, P<0.0001). Two-way ANOVA, followed by Dunnett’s multiple comparison test, individual P values of different time points versus before DT injection are labeled on the graph. (C) Although wave I amplitude change was randomly observed in control mice after DT injection, changes rarely persisted to 2 weeks after DT injection (n=10, P_16kHz_ <0.0001, P_24kHz_ <0.0001, P_32kHz-time_ <0.0001, p va). In contrast, significant reduction in wave I amplitude was observed in pericyte-depleted animals at 2 weeks after DT injection, particularly at high frequency (32 kHz) (n=10, P_8kHz_<0.0001, P_16kHz_ <0.0001, P_24kHz_ <0.0001, P_32kHz_ <0.0001). Two-way ANOVA, followed by Dunnett’s multiple comparison test, individual P values at 2 weeks after DT injection versus before DT injection at different sound pressure levels are labeled on the graph. Data are presented as the mean ± SEM.

In addition to the increased hearing threshold, we specifically examined ABR wave I, including its latency and amplitude, which is generally accepted as an indicator of auditory nerve activity in most mammalian animals and humans (Xie et al., 2018). We found the control mice showed no significant wave I latency change after DT injection. In contrast, the wave I latency in pericyte-depleted animals was significantly delayed, beginning at high frequency (32 kHz, after 4^th^ DT injection) and extending to lower frequency (8 kHz) at two weeks after DT injection (n=10, P<0.0001, two-way ANOVA) (**Fig. 3B**). Correspondingly, the significantly decreased amplitude of wave I were observed at all frequencies but started at much lower sound pressure level (dB) at high frequency (32 kHz) two weeks after DT injection (n=10, P<0.0001, two-way ANOVA) (**Fig. 3C**). Although a change in wave I amplitude was occasionally observed in control mice after DT injection, the change rarely persisted to two weeks after the DT injection.

In our pericyte-depleted animals, we observed significant SGN loss at all turns (**Fig. 4A and B** (n=9, P_Apex_=0.0085, P_Mid_=0.0099, P_Base_=0.0127, unpaired t-test) with markedly decreased β-III tubulin expression two weeks after DT treatment at the middle and basal turns (**Fig. 4A and C**, P_Apex_=0.1074, P_Mid_<0.0001, P_Base_<0.0001, unpaired t-test). β-III tubulin, a specific marker for neuronal cytoskeleton, is commonly used to distinguish between different types of neurons (Katsetos et al., 2003). Reduced tubulin is an indication of structural disassembly and it is seen in many neurodegenerative diseases as a sign of neuronal dysfunction (Baas et al., 2016). These results suggest loss of pericytes affects the viability of the spiral ganglion in adults.

**Figure 4.**
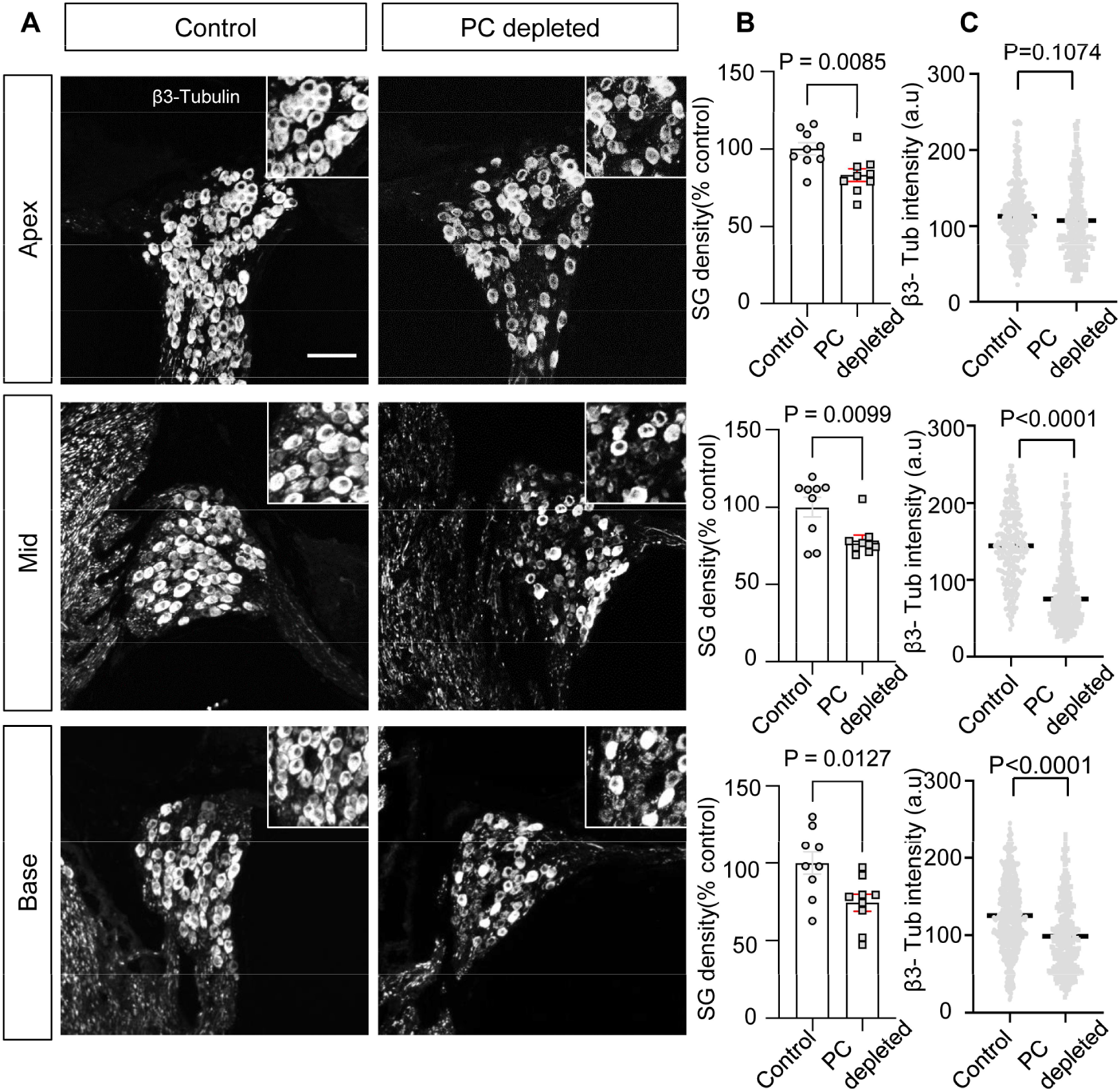
The depletion of pericytes led to SGN loss and decreased expression of β-III tubulin in SGNs. (A) Representative confocal images from control and pericyte-depleted animals, labeled with antibody for β-III tubulin. (B) Significant SGN loss at all turns two weeks after DT treatment (n=9, P_Apex_=0.0085, P_Mid_=0.0099, P_Base_=0.0127, unpaired t-test). (C) Significant decreased β-III tubulin expression in SGNs at middle and basal turns two weeks after DT treatment (P_Apex_=0.1074, P_Mid_<0.0001, P_Base_<0.0001, unpaired t-test). Data are presented as the mean ± SEM. Scale bar: D, 50 µm.

### Transcriptome analysis of cochlear pericytes

To gain a better understanding of the properties of cochlear pericytes, we investigated the transcriptome of primary cochlear pericytes cultured from postnatal (P10-P15) C57BL/6J mice. This cell line was generated from the stria vascularis by a well-established ‘mini-chip’ protocol as previously described (Neng et al., 2013), and passages 3-6 were used. Total RNA isolated from cultured cochlear pericytes was subjected to RNA-sequencing (RNA-seq) analysis. 12887 genes (RPKM>0) were identified expressed in mouse cochlear pericytes, 7889 (RPKM>0.5) of which were further analyzed by performing an overrepresentation test of Protein Analysis Through Evolutionary Relationships (PANTHER) pathways (relative to the whole-genome for Mus muscullus) in the PANTHER classification system (version 17.0) (Mi et al., 2013; Mi et al., 2019).

Finally, we identified 59 overrepresented PANTHER pathways in our cochlear pericyte dataset (**Table 1**), including angiogenesis related (“CCKR signaling map,” P06959; “Angiogenesis,” P00005; “VEGF signaling pathway,” P00056) and neurodegeneration related (“Alzheimer disease-presenilin pathway,” P00004; “Alzheimer disease-amyloid secretase pathway,” P00003; “5HT2 type receptor mediated signaling pathway,” P04374) pathways among the top 20 statistically significant results (**Fig. 5A**).

**Table 1.** Overrepresented PANTHER pathways for genes (RPKM>0.5) identified in cochlear pericytes.

| No. | PANTHER Pathways | % of gene in the list |  | -Log10 (FDR) |
| --- | --- | --- | --- | --- |
|  |  | Mus musculus (REF) | Pericyte |  |
| 1 | Inflammation mediated by chemokine and cytokine signaling pathway | 1.181979 | 5.072464 | 14.88941 |
| 2 | CCKR signaling map | 0.74101 | 3.804348 | 13.30277 |
| 3 | Angiogenesis | 0.813747 | 3.713768 | 11.61261 |
| 4 | Heterotrimeric G-protein signaling pathway-Gq alpha and Go alpha mediated pathway | 0.554621 | 2.717391 | 8.9914 |
| 5 | Heterotrimeric G-protein signaling pathway-Gi alpha and Gs alpha mediated pathway | 0.722826 | 2.98913 | 8.364516 |
| 6 | Wnt signaling pathway | 1.418375 | 4.257246 | 8.073658 |
| 7 | Gonadotropin-releasing hormone receptor pathway | 1.068327 | 3.623188 | 8.066513 |
| 8 | T cell activation | 0.409147 | 2.173913 | 7.876148 |
| 9 | Apoptosis signaling pathway | 0.563713 | 2.445652 | 7.314258 |
| 10 | Endothelin signaling pathway | 0.377324 | 1.992754 | 7.218245 |
| 11 | B cell activation | 0.322771 | 1.811594 | 6.931814 |
| 12 | VEGF signaling pathway | 0.304587 | 1.630435 | 5.958607 |
| 13 | Oxytocin receptor mediated signaling pathway | 0.272764 | 1.449275 | 5.242604 |
| 14 | Histamine H1 receptor mediated signaling pathway | 0.209119 | 1.268116 | 5.136677 |
| 15 | PI3 kinase pathway | 0.245488 | 1.358696 | 5.135489 |
| 16 | Thyrotropin-releasing hormone receptor signaling pathway | 0.286403 | 1.449275 | 5.086716 |
| 17 | Alzheimer disease-presenilin pathway | 0.577351 | 1.992754 | 4.640165 |
| 18 | Integrin signalling pathway | 0.863754 | 2.445652 | 4.315155 |
| 19 | Alzheimer disease-amyloid secretase pathway | 0.300041 | 1.358696 | 4.308919 |
| 20 | 5HT2 type receptor mediated signaling pathway | 0.309133 | 1.358696 | 4.194499 |
| 21 | Muscarinic acetylcholine receptor 1 and 3 signaling pathway | 0.272764 | 1.268116 | 4.154902 |
| 22 | PDGF signaling pathway | 0.650089 | 1.992754 | 4.017277 |
| 23 | Heterotrimeric G-protein signaling pathway-rod outer segment phototransduction | 0.177297 | 0.996377 | 3.886057 |
| 24 | EGF receptor signaling pathway | 0.618266 | 1.902174 | 3.882729 |
| 25 | Interleukin signaling pathway | 0.395508 | 1.449275 | 3.721246 |
| 26 | Muscarinic acetylcholine receptor 2 and 4 signaling pathway | 0.272764 | 1.177536 | 3.636388 |
|  |  | Mus musculus (REF) | Pericyte |  |
| 27 | FGF signaling pathway | 0.554621 | 1.721014 | 3.603801 |
| 28 | Cytoskeletal regulation by Rho GTPase | 0.363686 | 1.358696 | 3.595166 |
| 29 | Nicotinic acetylcholine receptor signaling pathway | 0.450061 | 1.449275 | 3.19382 |
| 30 | p53 pathway feedback loops 2 | 0.227304 | 0.996377 | 3.157391 |
| 31 | ATP synthesis | 0.031823 | 0.452899 | 3.123782 |
| 32 | Axon guidance mediated by netrin | 0.159113 | 0.815217 | 3.010105 |
| 33 | Beta1 adrenergic receptor signaling pathway | 0.209119 | 0.905797 | 2.886057 |
| 34 | Enkephalin release | 0.168205 | 0.815217 | 2.882729 |
| 35 | GABA-B receptor II signaling | 0.168205 | 0.815217 | 2.869666 |
| 36 | Metabotropic glutamate receptor group II pathway | 0.213665 | 0.905797 | 2.832683 |
| 37 | Insulin/IGF pathway-protein kinase B signaling cascade | 0.177297 | 0.815217 | 2.772113 |
| 38 | Blood coagulation | 0.236396 | 0.905797 | 2.551294 |
| 39 | FAS signaling pathway | 0.154567 | 0.724638 | 2.522879 |
| 40 | Nicotine pharmacodynamics pathway | 0.159113 | 0.724638 | 2.462181 |
| 41 | Dopamine receptor mediated signaling pathway | 0.25458 | 0.905797 | 2.376751 |
| 42 | Beta2 adrenergic receptor signaling pathway | 0.209119 | 0.815217 | 2.366532 |
| 43 | Alpha adrenergic receptor signaling pathway | 0.10456 | 0.543478 | 2.095284 |
| 44 | Axon guidance mediated by Slit/Robo | 0.113652 | 0.543478 | 1.978811 |
| 45 | TGF-beta signaling pathway | 0.459154 | 1.177536 | 1.978811 |
| 46 | p53 pathway | 0.400055 | 1.086957 | 1.974694 |
| 47 | Androgen/estrogene/progesterone biosynthesis | 0.077283 | 0.452899 | 1.970616 |
| 48 | Endogenous cannabinoid signaling | 0.113652 | 0.543478 | 1.970616 |
| 49 | Corticotropin releasing factor receptor signaling pathway | 0.154567 | 0.634058 | 1.970616 |
| 50 | Histamine H2 receptor mediated signaling pathway | 0.118198 | 0.543478 | 1.931814 |
| 51 | 5HT1 type receptor mediated signaling pathway | 0.209119 | 0.724638 | 1.910095 |
| 52 | Opioid proopiomelanocortin pathway | 0.168205 | 0.634058 | 1.853872 |
| 53 | Hypoxia response via HIF activation | 0.12729 | 0.543478 | 1.821023 |
| 54 | TCA cycle | 0.054553 | 0.362319 | 1.754487 |
| 55 | Cadherin signaling pathway | 0.736464 | 1.539855 | 1.686133 |
| 56 | Pyruvate metabolism | 0.059099 | 0.362319 | 1.669586 |
| 57 | Triacylglycerol metabolism | 0.009092 | 0.181159 | 1.446117 |
| 58 | Opioid prodynorphin pathway | 0.163659 | 0.543478 | 1.407823 |
| 59 | Opioid proenkephalin pathway | 0.172751 | 0.543478 | 1.321482 |

**Table 1-Source data 1. Gene list of cochlear pericytes identified by RNA-seq analysis.**

**Figure 5.**
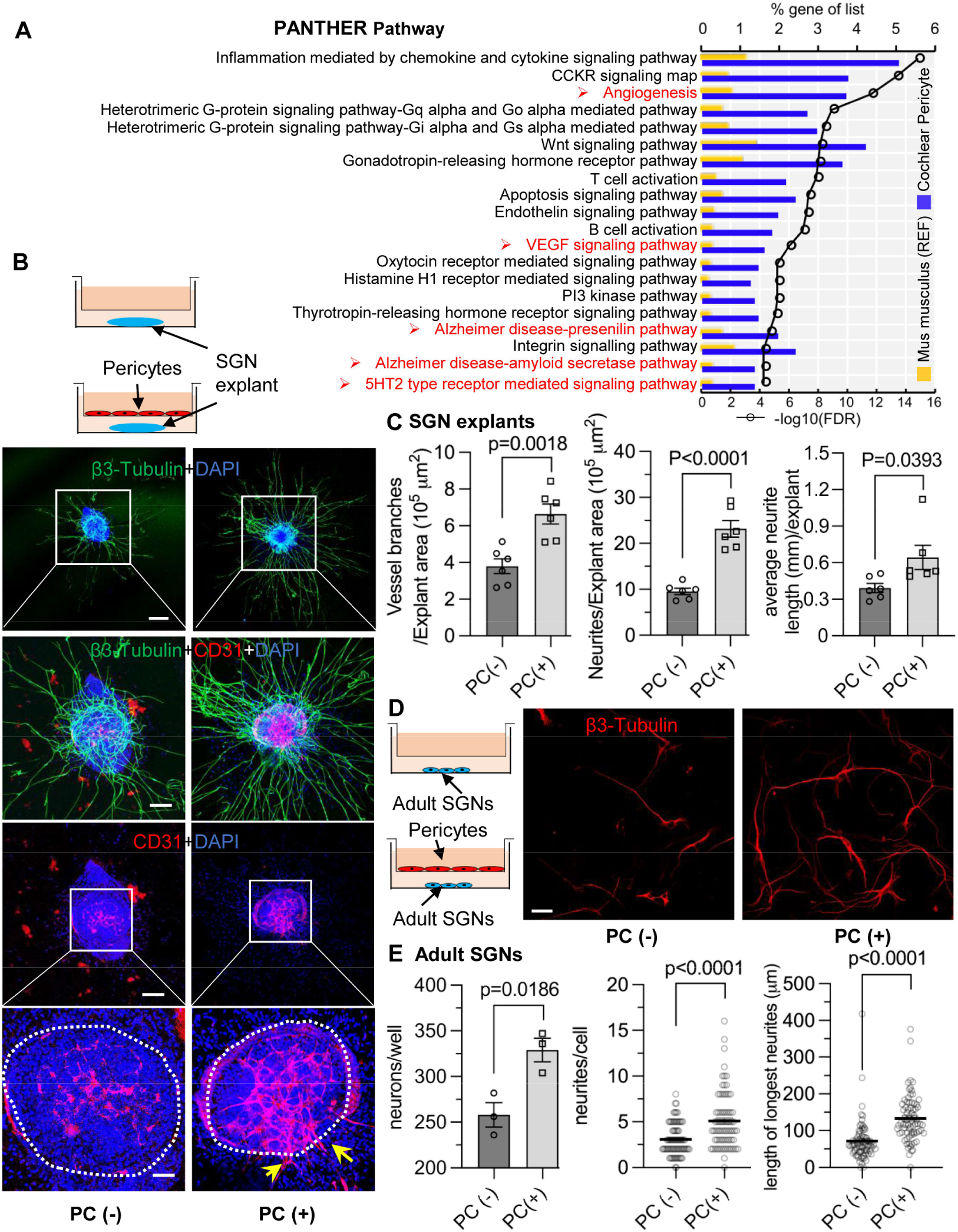
Pericytes promote both vascular and neuronal growth in spiral ganglion *in vitro*. (A) The 20 most significant overrepresented PANTHER pathways for genes (RPKM>0.5) identified in cochlear pericytes. (B) Neonatal SGN explants co-cultured with pericytes show robust SGN dendritic growth (green, labeled with β-III Tubulin) and new vessel growth (red, labeled with CD31). (C) There are significant differences in number of new vessel branches and in number and length of dendritic fibers in the two groups (n=6, P_Vascular branch #/area_ =0.0018, P_Neurites #/area_ <0.0001, P_Neurite length/explant_ =0.0393, unpaired t test). (D) Adult SGNs co-cultured with pericytes show robust SGN dendritic growth (red, labeled with β-III Tubulin). (E) There are significant differences in cell survival, and in average neurite number and length, in the two groups (n=3 wells per group, 25 cells per well, P_neuron survival_ =0.0186, P_Neurites #/cell_ <0.0001, P_longest neurite length/cell_ <0.001, unpaired t test). Data are presented as the mean ± SEM. Scale bars: B, 300 µm (top), 150 µm (middle), 50 µm (bottom); D, 50 µm.

### Pericytes promote vascular and neuronal growth in the spiral ganglion *in vitro*

We next investigated whether pericytes directly communicate with nearby cells and affect the corresponding vascular and neuronal biology. Neonatal SGN explants and adult SGNs were cultured with exogenous pericytes in a transwell co-culture system, as illustrated in **Figs. 5B and D**. After five days in culture, we observed numerous and longer dendritic fibers, as well as new vascular sprouting, in the exogenous pericyte treated SGN explants relative to controls (**Fig. 5B**). There were significant differences in the number of new vessel branches, and in number and length of dendritic fibers between the two groups (n=6, P_Vascular branch#/area_=0.0018, P_Neurites #/area_<0.0001, P_Neurite length/explant_ =0.0393, unpaired t test) (**Fig. 5C**). The exogenous pericytes also consistently promoted survival and new neurite growth in adult SGNs (**Fig. 5D**). Significant differences in cell survival, and in average neurite number and length, were observed between the two groups (n=3 wells, 25 cells per well, P_neuron survival_=0.0186, P_Neurites#/cell_<0.0001, P_longest neurite length/cell_<0.0001, unpaired t test) (**Fig. 5E**). The data clearly indicate that pericytes mediate both vascular and neural growth through extracellular communication in both neonatal and adult SGN tissue, as well as demonstrate that pericytes are essential for vascular and neuronal growth both during development and in the adult.

### Identification of cochlear pericyte-derived exosomes

We then asked how pericytes communicate with neighboring cells. Many cells secrete exosomes, carrying cargo including all known molecular constituents of the host cell, including protein, DNA, RNA, lipids, and metabolites. These are transported to surrounding cells (Bang & Thum, 2012) to effect biological function in the receiving cells (Dai et al., 2020; Kalluri & LeBleu, 2020). To determine whether cochlear pericyte-derived exosomes contribute to the regulation of vascular and neuronal growth in spiral ganglia, we first explored the properties of EVs isolated from pericyte-conditioned culture media. EVs were purified from the collected media by ultrafiltration and size exclusion separation, as illustrated in **Fig. 6A**. Using nanoparticle tracking analysis (NTA), a widely employed technique for the characterization of particles in liquids (Dragovic et al., 2011), we found a rich population of exosome-sized (∼50–150 nm) particles, 96.4% of which are labeled with Exoglow (a membrane EV marker kit), as shown in **Fig. 6B**. Further transmission electron microscopy (TEM) images showed a classic cup-shaped structure in these membrane vesicles, which is consistent with previously described exosomes from other cell types, such as HCT116 cells (Jung & Mun, 2018) and utricles (Breglio et al., 2020). Moreover, proteomic analysis of purified pericyte-derived exosomes identified 580 proteins, 496 of which overlapped with genes that we identified in cochlear pericytes (**Fig. 6C**), including the common exosome markers CD9, CD63, CD81 and Tsg101, as shown in **Table 2**. Together, these results indicate the pericytes constitutively release exosomes.

**Table 2.** Common protein markers for exosomes were identified in the pericyte-derived exosomes.

| Protein | Gene names | # PSMs | Log2 LFQ intensity |
| --- | --- | --- | --- |
| CD81 antigen | *Cd81* | 26 | 29.6153 |
| CD63 antigen | *Cd63* | 3 | 26.1908 |
| CD9 antigen | *Cd9* | 16 | 29.2576 |
| Tumor susceptibility gene 101 | *Tsg101* | 5 | 25.7598 |

**Figure 6.**
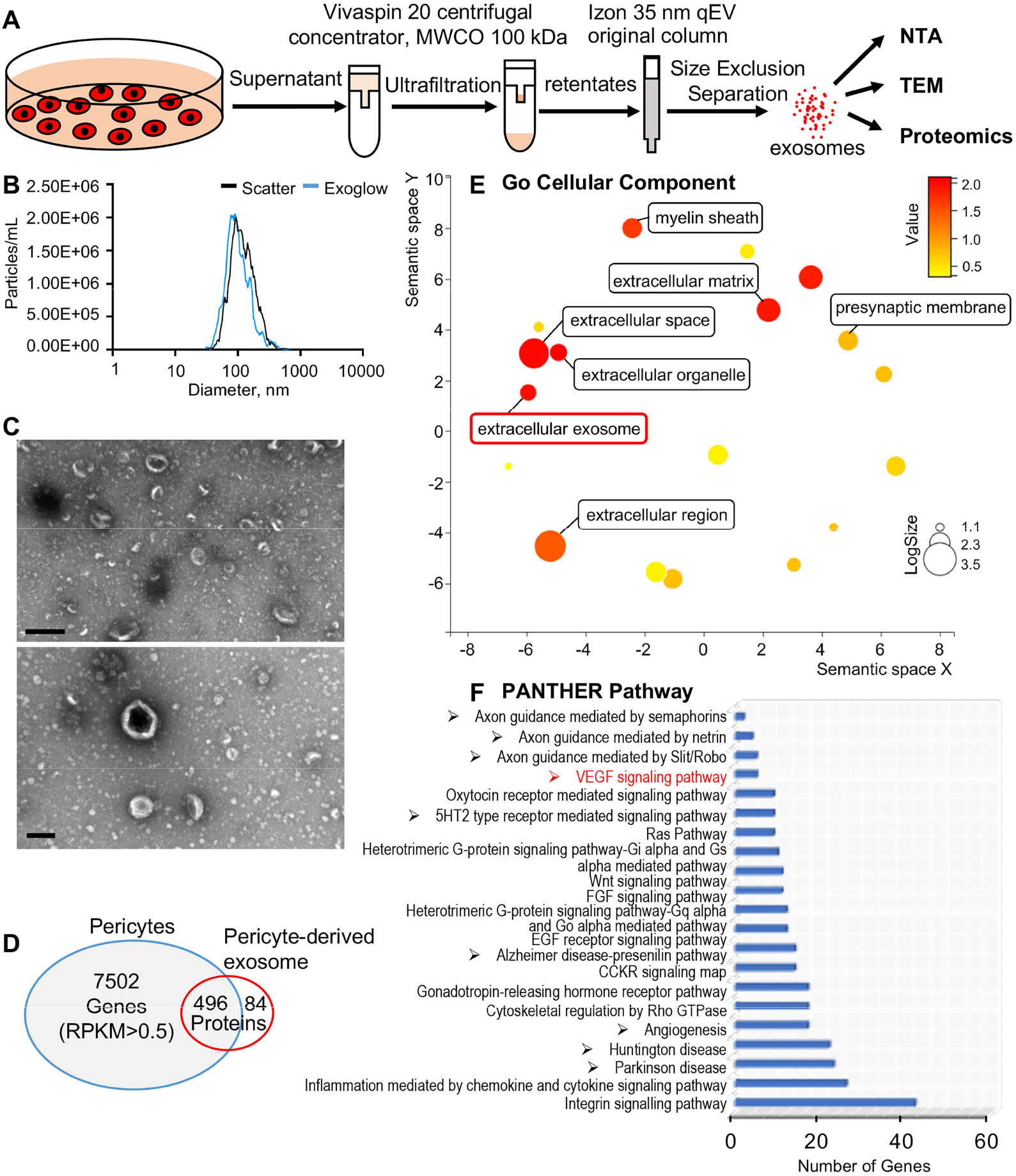
Identification of pericyte-derived exosomes. (A) Schematic of the exosome purification procedure used to isolate exosomes from pericyte-conditioned culture medium for NTA, TEM, and proteomics analysis. Exosomes were isolated via ultracentrifugation followed by size-exclusion separation. (B) NTA shows a rich population of exosome-sized (∼50–150 nm) particles, 96.4% of which are labeled with Exoglow (a membrane EV marker kit). (C) TEM images show a classic cup-shaped structure membrane vesicles with diameter around 100 nm. (D) Proteomics analysis of exosomes identified a total of 570 unique protein families in exosomes, 496 of which overlap with genes identified in pericyte isolated total RNA. (E) The 22 most significantly enriched GO cellular component terms for proteins identified in the pericyte-derived exosomes. (F) The relative PANTHER pathways identified in pericyte-derived exosome proteins. Scale bars: C, 500 nm (top), 200 nm (bottom). **Figure 6-Source data 1. Protein families of cochlear pericyte-derived exosomes identified by proteomics analysis**.

We ran Gene Ontology (GO) enrichment analysis on these proteins using the PANTHER classification system, followed by removal of redundant ontology terms using Reduce and Visualize Gene Ontology (REVIGO) (Supek et al., 2011). Analysis by cellular component class identified enrichment of 22 GO terms, including “extracellular exosomes” (GO: 0070062), neurotransmission related terms such as “myelin sheath” (GO: 0043209) and “presynaptic membrane” (GO: 0042734) (**Fig. 6D**). Although analysis by PANTHER pathways didn’t identify significantly enriched pathways, we listed the identified pathways in the order of relative protein number (**Fig. 6E**) and found that the angiogenesis related (“Angiogenesis,” P00005; “VEGF signaling pathway,” P00056) and neurodegeneration related (“Parkinson disease,” P00049; “Huntington disease,”P00029) pathways were among the top ranks. These pathways are consistent with pathways we identified in cochlear pericytes, as shown in **Fig. 5A**. Collectively, these data strongly indicate that cochlear pericyte-derived exosomes participate in regulation of vascular and neuronal growth in the spiral ganglion.

### Cochlear pericyte-derived exosomes promote vascular and neuronal growth via VEGFR2 signaling

VEGF/VEGFR2 signaling plays a key role in formation and growth of blood vessels, but is also implicated in neurodegeneration (Storkebaum & Carmeliet, 2004). Our bulk RNA-seq analysis identified VEGF-A expression in cochlea pericytes, as did RT-PCR (**Fig. 7A**). Although VEGF-C was also in the list, the expression was very low (RPKM=0.07301). ELISA protein analysis showed 3.696 ± 0.003 ng of VEGF-A/ 8 × 10^5^ pericytes released into 4 ml of culture medium without growth factors after 3 days culture (**Fig. 7B**). In contrast, when we depleted the pericytes in vivo, we found significantly less production of VEGF-A in the cochlea (**Fig. 7C**).

**Figure 7.**
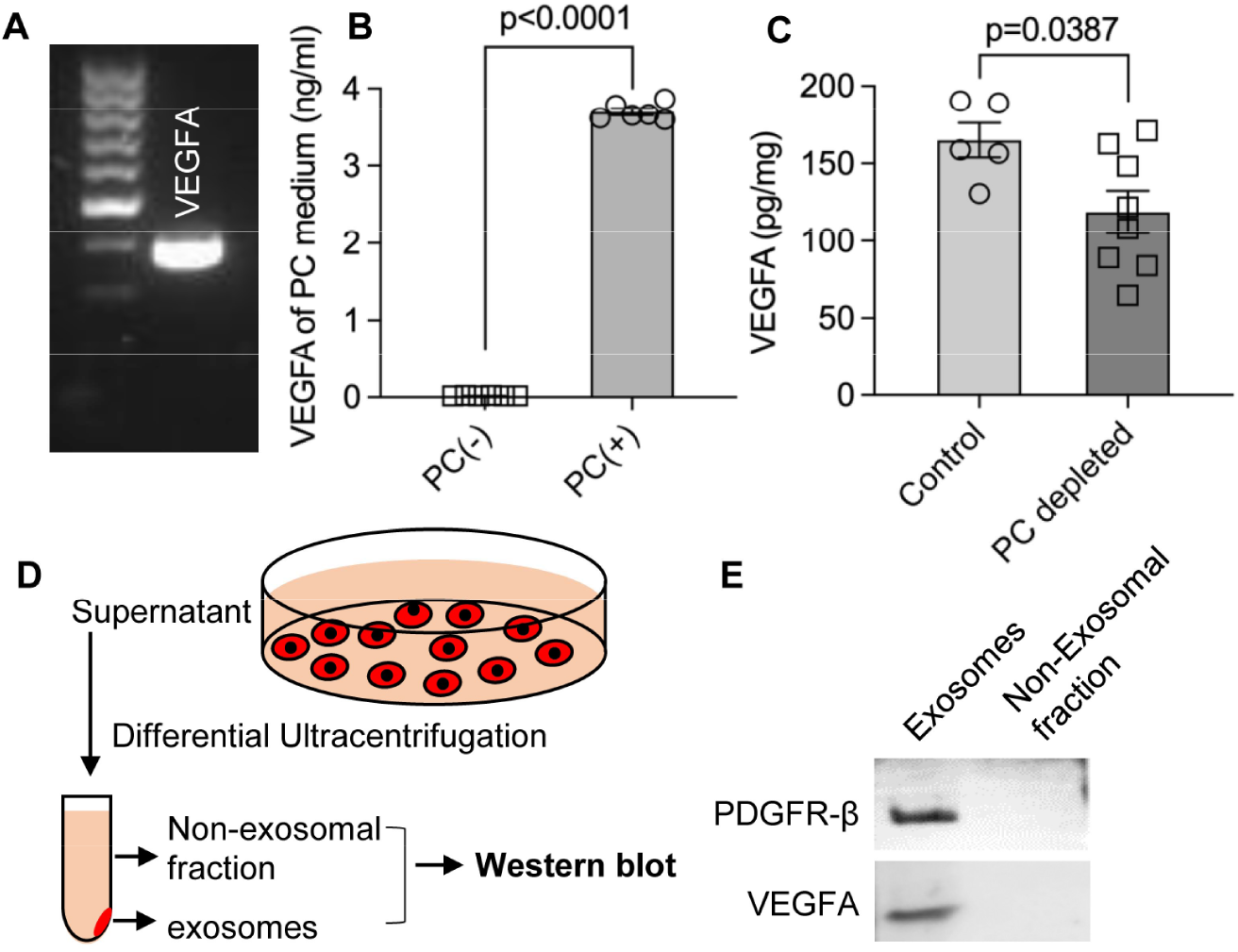
Pericytes release VEGFA through exosomes. (A) mRNA expression of VEGF-A in primary cochlear pericytes. (B) VEGF-A production assessed by ELISA at day 3 in the control and pericyte containing culture medium (n=6, P <0.0001, unpaired t test). (C) VEGFA expression level assessed by ELISA in the cochlea of control and pericyte-depleted mice (n_control_=5, n_pericyte depletion_=8, P=0.0387, unpaired t test). (D) Exosomes and the non-exosomal fraction (supernatant) were purified from pericyte-conditioned media using differential ultracentrifugation for Western blot analysis. (E) Western blot showing the expression of VEGFA in exosomes. Similarly, PDGFR-β (pericyte membrane marker) was detected exclusively in exosomes. Data are presented as the mean ± SEM. **Figure 7-Source data 1. original uncropped blots of VEGFA and PDGFR-β, and raw gel of whole protein staining with the relevant bands clearly labelled**.

Is VEGF-A largely transported by pericyte-derived exosomes or is it directly secreted into the medium? To determine the answer, the exosome and non-exosomal supernatant fraction of the pericyte culture medium were isolated by differential ultracentrifugation, and analyzed by Western blot (**Fig. 7D**). We found that both PDGFRβ (a pericyte cellular membrane marker) and VEGF-A were expressed in the exosomes, but not detected in the non-exosomal supernatant fraction (**Fig. 7E**). We next asked about the receptor(s) through which VEGF-A exerts its effect on vascular and neuronal growth. Various neural cells express one or more of the known VEGF receptors and could thus directly respond to VEGF released by neighboring cells (D’Amore, 2007; Ogunshola et al., 2002; Teran & Nugent, 2019). VEGFR2 is the major VEGF receptor with a critical role in angiogenesis (Abhinand et al., 2016). It also has been reported to mediate the VEGF signaling in neurons, with roles in neurogenesis, neuronal survival, and axonal growth (Bellon et al., 2010; Luck et al., 2019). We used immunofluorescence to identify VEGFR2 expression in both SGNs and blood vessels, as shown in **Fig. 8A-D**. Our findings lead us to hypothesize that pericyte-derived exosomes promote vascular and neuronal growth in spiral ganglia through a VEGFR2 signaling pathway.

**Figure 8.**
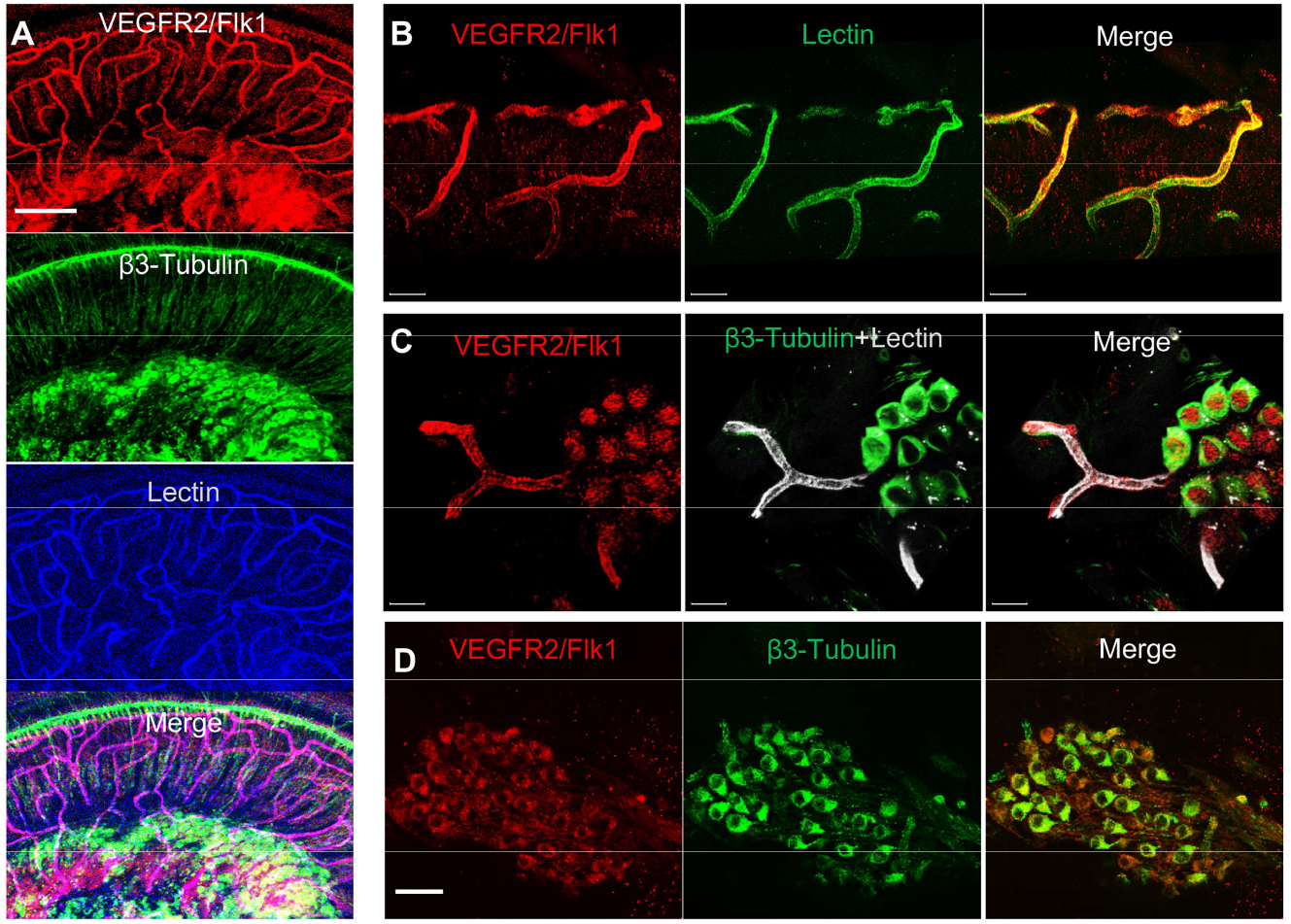
VEGFR2 expression in the spiral ganglion region. (A-C) Representative confocal images of a cochlear whole mount under low (A) and high (B and C) magnification. (D) Cross section showing VEGFR2 (red) is positively expressed in both SGNs (labeled for β-III Tubulin) and blood vessels (labeled for lectin). Scale bars: E, 100 µm; F, 30 µm; G, 20 µm; H, 50 µm.

To test our hypothesis, we directly treated neonatal SGN explants and adult SGNs with purified exosomes isolated with total exosome isolation reagent. We found the exosome treatment promoted both vascular and neuronal growth in neonatal SGN explants, which was attenuated by the specific VEGFR2 inhibitor, SU5408 (Roskoski, 2017) at 100 nM (n=6, P_Vascular branch #/area_<0.0001; P_Neurites #/area_<0.0001; P_Neurites #/area_=0.0268, one-way ANOVA) (**Fig. 9B and D**). Similarly, exosome treatment also promoted the survival and neurite growth of adult SGNs, which was also arrested by SU5408 (n=4 wells, 25 cells per well, P_cell survial_=0.0048, P_Neurites #/cell_<0.0001, P_longest neurite length/cell_ <0.0001, one-way ANOVA) (**Fig. 9C and E**). The results indicate pericyte-derived exosomes promote vascular and neuronal growth through a VEGFA/VEGFR2 signaling pathway. Dose-response of SU5408 was assessed in a co-cultured of pericytes and neonatal SGN explants (**Fig. 10A-C**).

**Figure 9.**
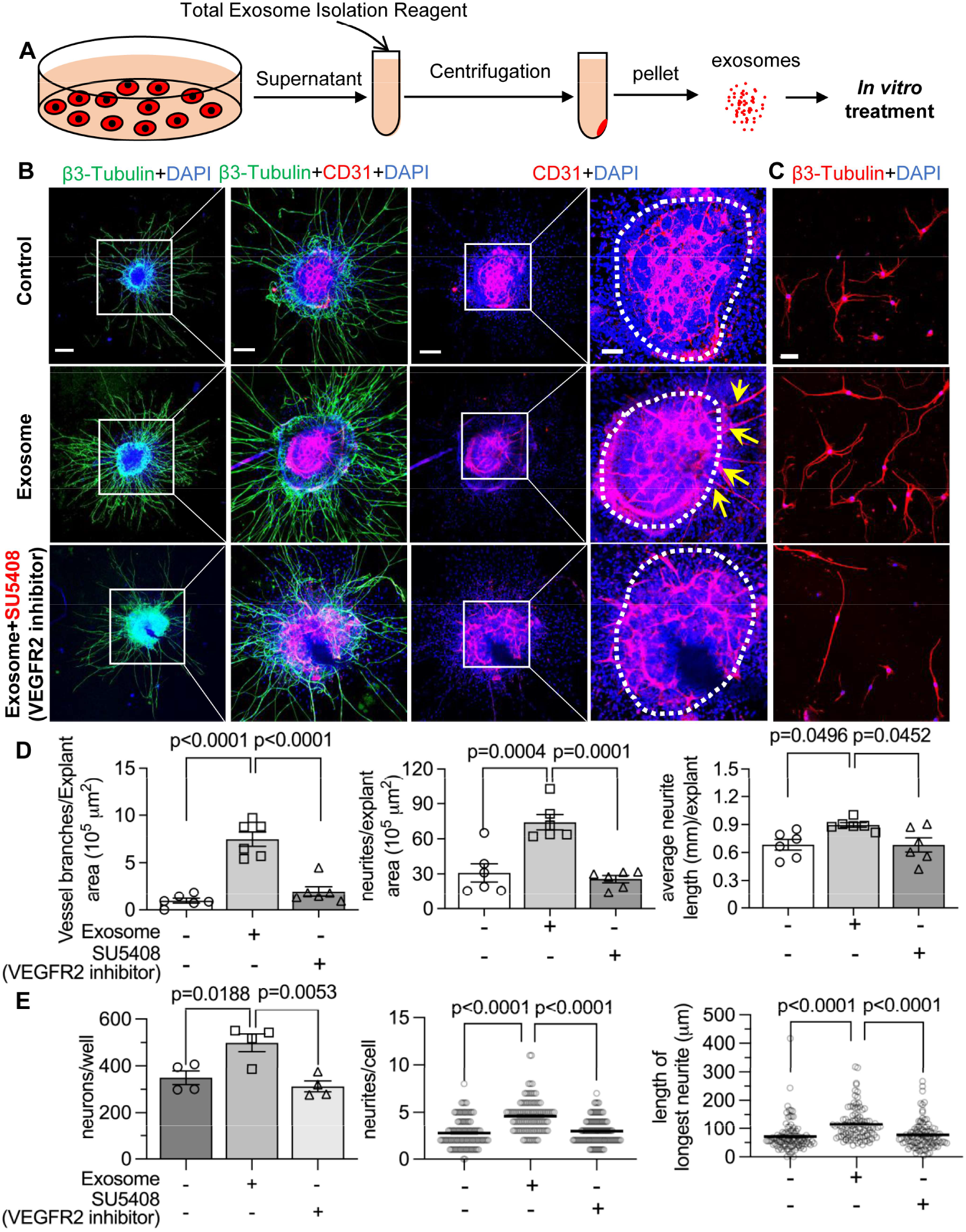
Pericyte-derived exosomes promote angiogenesis and SGN growth through VEGFR2 signaling. (A) Exosomes was purified from pericyte-conditioned media using total exosome isolation reagent for *in vitro* treatment. (B) Compared to the control group, exosome treated SGN explants showed robust SGN dendritic growth (green, labeled with β-III Tubulin) and new vessel growth (red, labeled for CD31). In contrast, both neurogenic and angiogenetic activity were decreased when a VEGFR2 inhibitor, SU5408, was presented in the medium. (C) Exosome treated adult SGNs showed more SGN dendritic growth (red, labeled for β-III Tubulin) compared with control and VEGFR2 inhibitor groups. (D) There are significant differences in new vessel branch number and in dendritic fiber number and length in the three groups (n=6, P=0.0017). (E) There are significant differences in cell survival, and in average neurite number and length, in the three groups (n=4 wells per group, 25 cells per well, P=0.0048). One-way ANOVA followed by Tukey’s multiple comparison test, individual P values of different group comparisons are labeled on the graph. Data are presented as the mean ± SEM. Scale bars: B, 300 µm (left), 150 µm (center), 50 µm (right); C, 50 µm.

**Figure 10.**
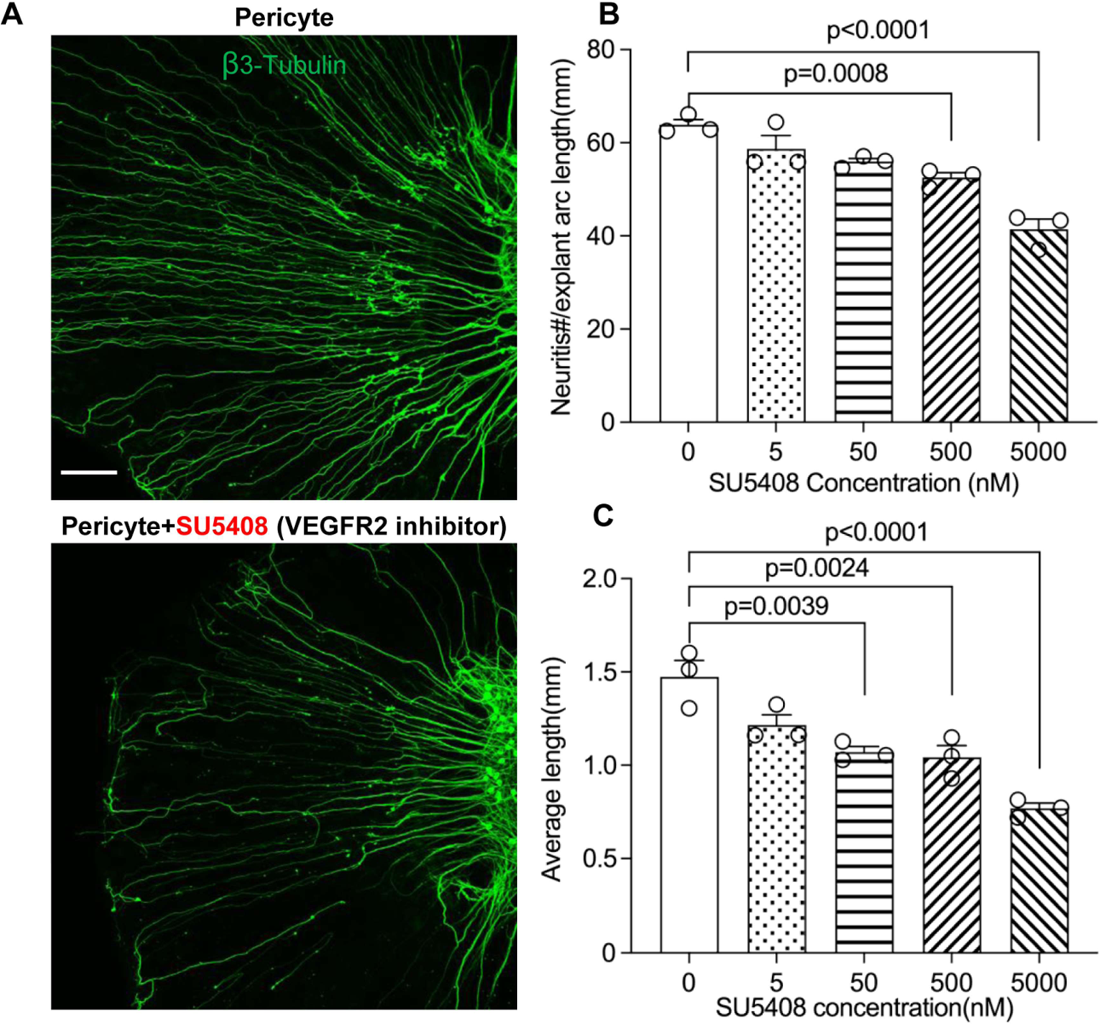
VEGFR2 blockage causes reduction of neuronal dendritic growth is seen in pericytes are co-cultured SGN explants. (A) Representative images showing the pattern of SGN dendritic growth under different experimental conditions. (B and C) Number of dendritic fibers and length is inhibited in a dose-dependent manner (n= 3, P_number_ < 0.0001, P_length_ < 0.0001, one-way ANOVA followed by a Tukey’s multiple comparison test, individual P values between different groups are labeled on the graph). Data are presented as the mean ± SEM. Scale bar: A, 150 µm.

## Discussion

Pericytes are prevalent on microvessels in the cochlea, but they have not received much research attention. In this investigation, we used an inducible and conditional pericyte depletion mouse model (*Pdgfrb*^*CreERT2*^; *R26*^*iDTR*^) to demonstrate that pericytes in the adult ear are critical for the stability of mature vessel beds and viability of SGNs in the region of the peripheral spiral limbus. Depletion of pericytes causes loss of vascular volume and SGNs, with concomitant loss of hearing sensitivity. We report on a pericyte-released growth factor, VEGF-A, conveyed by exosomes, shown to strongly mediate and promote SGN survival and growth by binding to VEGFR2 (Flk-1). This study provides the first clear evidence that pericytes have a critical role in vascular and neuronal health in the adult ear. Without the normal disposition of pericytes, animals lose hearing

Recent research has highlighted the extent to which pericytes play a critical role in vascular function and neuronal protection in the central nervous system (CNS) (Bergers & Song, 2005; Brown et al., 2019). Studies, however, on the role of pericytes in the peripheral nervous system are limited. In the CNS, loss of pericytes causes brain infarction with diminished capillary perfusion and blood flow (Armulik et al., 2010; Bell et al., 2010). In the mouse brain, damaged pericytes effectively constrict capillaries and induce a long-lasting reduction in cerebral blood flow, even after recanalization of the larger vessels, and it causes a no-reflow condition (Hall et al., 2014; Kloner et al., 2018; Yemisci et al., 2009). Loss of pericytes also interrupts blood-brain-barrier integrity (Armulik et al., 2010; Bell et al., 2010; Nikolakopoulou et al., 2019). In peripheral organs such as the ear, we previously reported that pericytes control perfusion of cochlear blood flow (Dai et al., 2010). Loss of pericytes leads to increased cochlear permeability and reduced endocochlear potential (Zhang et al., 2021). The results obtained in this study further demonstrate the important role of pericytes in maintaining vascular stability in peripheral systems such as auditory SGNs in the cochlea.

SGNs are neural elements transmitting localized electrical activity from hair cells to second-order neurons in the cochlear nucleus (Coate et al., 2019). The SGNs are vulnerable to interruption in blood supply since the energy demand of the SGNs is directly supplied by the surrounding blood circulation (Jiang et al., 2019; Mei et al., 2020). The microvascular network in the region is densely populated by pericytes (Jiang et al., 2019). Loss of pericytes affects SGN viability and causes significant loss of SGNs across all turns as shown in **Fig. 4**. β-III tubulin, a specific marker for neuronal cytoskeleton, is commonly used to distinguish neurons from other cell types (Katsetos et al., 2003). We found significantly decreased expression of β-III tubulin protein in SGNs in pericyte-depleted animals, particularly at the middle and basal turns.

Tubulin is the primary structural protein in microtubules, and the assembly and stability of tubulin are essential to the structure and function of neurons. A gradual loss of neuronal microtubule mass is seen in many neuro-degenerative diseases (Baas et al., 2016). ABR functional studies show a delay in latency and low amplitude starts at the basal turn and moves to middle and low range frequencies. The mechanism underlying the vulnerability of SGNs at the basal turn is not known. However, the significantly decreased expression of β-III tubulin protein in SGNs at the basal and middle turns may indicate that SGN damage is more severe than at the apical turn. In addition, a recent single-cell analysis study of mouse SGNs showed there were three functionally distinct subtypes of SGN, each displaying molecular variation along the tonotopic axis (Shrestha et al., 2018). Sensitivity at the different turns to pericyte loss could also be due to SGN diversity and the variance in SGN vulnerability to environmental perturbation. Further study is needed to define the mechanisms of SGN vulnerability.

What is the underlying mechanism of pericyte-loss-related damage to SGNs? Our data suggest both blood flow dysfunction and direct disruption of pericyte-SGN communication are at issue. It is well-documented that pericyte loss triggers primary vascular dysfunction, leading to neurodegeneration (Winkler et al., 2011). For example, in the CNS, pericyte loss induces vascular leakage, insufficient blood perfusion, and tissue hypoxia, ultimately leading to neuron degeneration (Quaegebeur et al., 2010; Sagare et al., 2013; Winkler et al., 2011; Zlokovic, 2011). We previously demonstrated active communication between pericytes, endothelial cells, and SGNs in the cochlea (Jiang et al., 2019). SGN uptake of pericyte-released particles *in vivo* was consistently seen in the study, as shown in **Fig. 11**. Pericytes release many growth factors relevant to maintenance of organ homeostasis (A. Gaceb, M. Barbariga, et al., 2018; A. Gaceb, I. Ozen, et al., 2018; Gaceb & Paul, 2018). In this study, we used bulk-RNA-seq analysis on the purified cochlear pericytes to identify the different vascular- and neuronal-growth factors in our mouse cochlear pericyte dataset. We found pericyte-related angiogenesis and neuroprotective pathways overrepresented (**Fig. 5A**). In pericyte co-culture models of neonatal SGN explants and adult SGNs, we found pericytes to induce robust vascular and neuronal dendritic growth in the neonatal SGN explants and to increase cell survival and neurite growth in the adult SGNs, as shown in **Fig. 5B-E**. The results suggest direct intercellular communication between pericytes and SGNs in addition to direct effects exerted by pericytes on vascular function.

**Figure 11.**
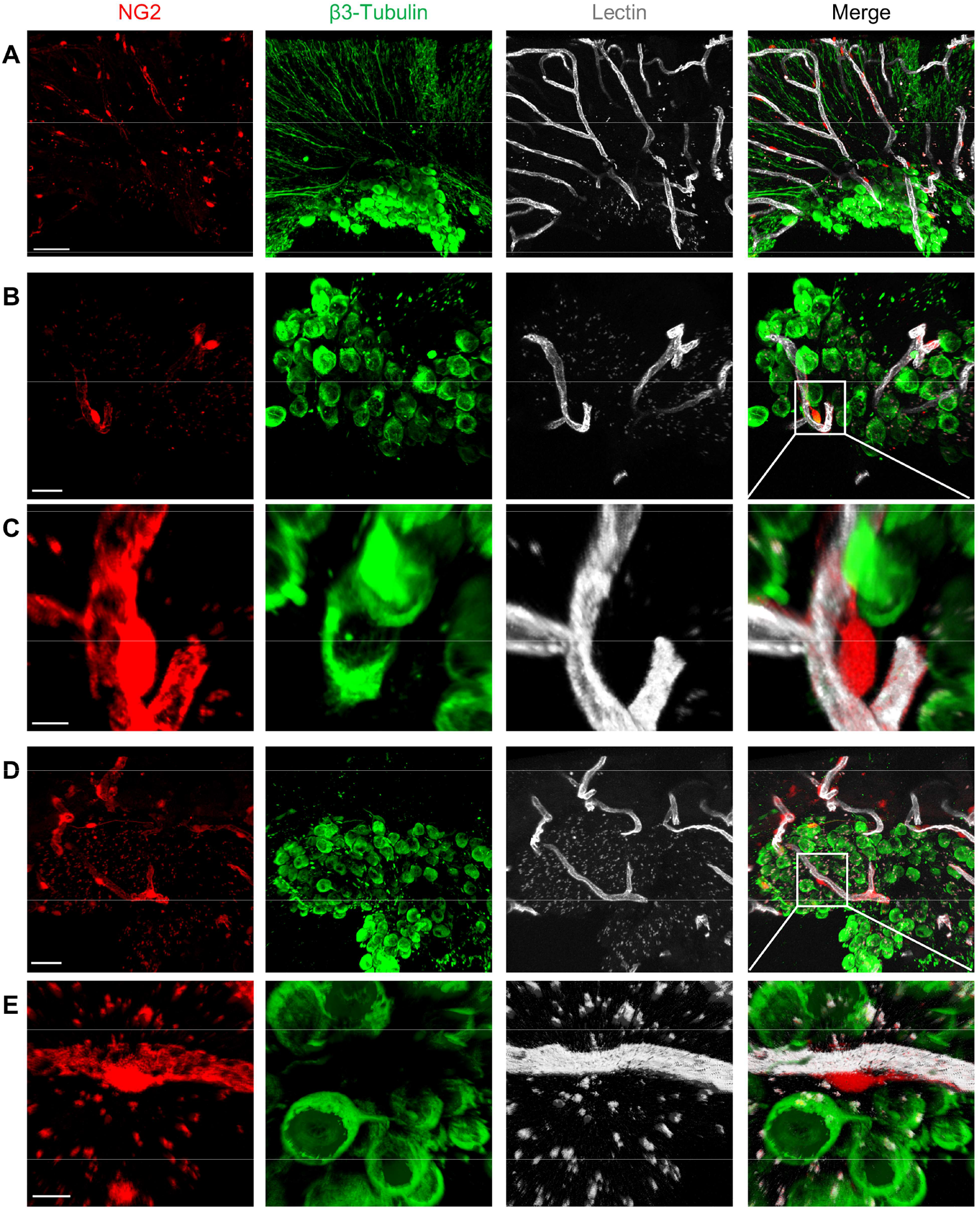
Cochlear vascular networks in the region of the spiral limbus are densely populated by PCs, and SGNs take up particles released by the pericytes. (A-C) Confocal projection images of the spiral limbus from an *NG2*^*DsRedBAC*^ mouse showing pericytes (red) situated on microvessels labeled with Lectin-Alexa Fluor® 649, surrounding the SGNs under low and high magnifications. The peripheral fibers are labeled with an antibody for β-III tubulin. (D and E) Confocal projection images showing the cross-talk between SGNs and PCs indicated by red fluorescent particles (red) taken up by SGNs (green). Scale bars: A, 50 µm; B, 20 µm; C, 5 µm; D, 40 µm; E, 5 µm.

How do pericytes communicate with endothelial cells and SGNs? Many cells release exosomes, nano-sized extracellular vesicles (50-150 nm diameter), which transport cargo such as nucleic acids, proteins, and lipids. These exosomes have significant physiological effects in the recipient cells (Bang & Thum, 2012; Colombo et al., 2014; Zhang et al., 2019). Pericyte-released exosomes were first reported by Gaceb et al. in 2018 (Abderahim Gaceb et al., 2018). Pericyte-derived exosome physiology, including characterization of size and morphology, remains limited. Recent studies have demonstrated that pericyte-derived exosomes participate in regulation of microvascular function under different pathological conditions (Ye et al., 2021; Yuan et al., 2019). In our study the exosomes displayed the classic cup-shaped morphology (**Fig. 6B and C)**. Further proteomic analysis of these particles identified 580 proteins enriched in the GO of cellular components in extracellular exosomes, including proteins relating to angiogenesis, neurodegeneration, and neuroprotection (**Fig. 6E-F**). Our findings are strong indication of participation of cochlear pericyte-derived exosomes in intercellular communication with spiral ganglion.

How do pericyte-derived exosomes affect angiogenesis and neurogenesis? VEGF/VEGFR2 signaling is a well-defined, classical pathway that plays a critical role in both angiogenesis and neuroprotection (Storkebaum & Carmeliet, 2004). VEGFR2 is typically expressed in both vascular cells and SGNs (**Figs. 8A-D**). In this study, we used *in vitro* neonatal SGN explant and adult SGN tissue culture models to specifically investigate the VEGF-A controlled angiogenesis signaling pathway in relation to vascular and neuronal growth. Our results show blockage of the VEGFR2 receptor with a specific VEGFR2 inhibitor, SU5408 (Roskoski, 2017), dramatically halts angiogenesis and neuronal growth (**Fig. 9**), indicating that pericyte-derived VEGF-A-containing exosomes promote both vascular and neuronal growth through a VGFR2 signaling pathway. The exact mechanism of the VEGFR2 regulation and downstream signaling remains unclear. Nevertheless, the data clearly indicate cochlear angiogenesis and neurogenesis require VEGF-A signaling. A recent study by Okabe et al. reports that neuron-derived VEGF contributes to cortical and hippocampal development, but is independent of VEGFR1/2-mediated neurotropism (Okabe et al., 2020). They conclude that neuron-derived VEGF contributes to cortical and hippocampal development likely through angiogenesis independent of any direct neurotrophic effects mediated by VEGFR1 and 2. Although there is still no consensus on whether cochlear angiogenesis affects auditory peripheral neurogenesis, our co-culture model of pericytes and adult SGNs does provide evidence the VEGF-A/VEGFR2 signal strongly promotes auditory SGN dendritic growth, as demonstrated in **Fig. 9C**. In addition, cochlear pericyte derived-exosomes contain numerous other proteins, as we identified in our proteomic analysis. For example, they contain messenger RNA and micro-RNAs not investigated in this study. Some of these may also be playing an important role in regulation of angiogenesis, SGN development, and neurite growth. However, it must be emphasized that treatment with the VEGFR2 inhibitor almost completely blocked the effect of the exosomes relative to control groups (see **Fig. 9**). This suggests the VEGF-A/VEGFR2 signaling is the dominant factor regulating vascular and neuronal growth. However, this does not obviate the possibility the underlying mechanism of pericyte-related neuroprotection is more complicated than what we have discovered in this study.

In summary, our data provide the first clear-cut experimental evidence that pericytes are essential for the maintenance of cochlear vascular stability and SGN viability in adults. Loss of pericytes disrupts vascular structure and damages SGNs. The damage is related to the impairment of VEGF-A mediated communication between pericytes, endothelial cells, and SGNs, as illustrated in **Fig. 12**. These findings demonstrate for the first time that pericyte-released exosomes have a major role in vascular and neuronal health.

**Figure 12.**
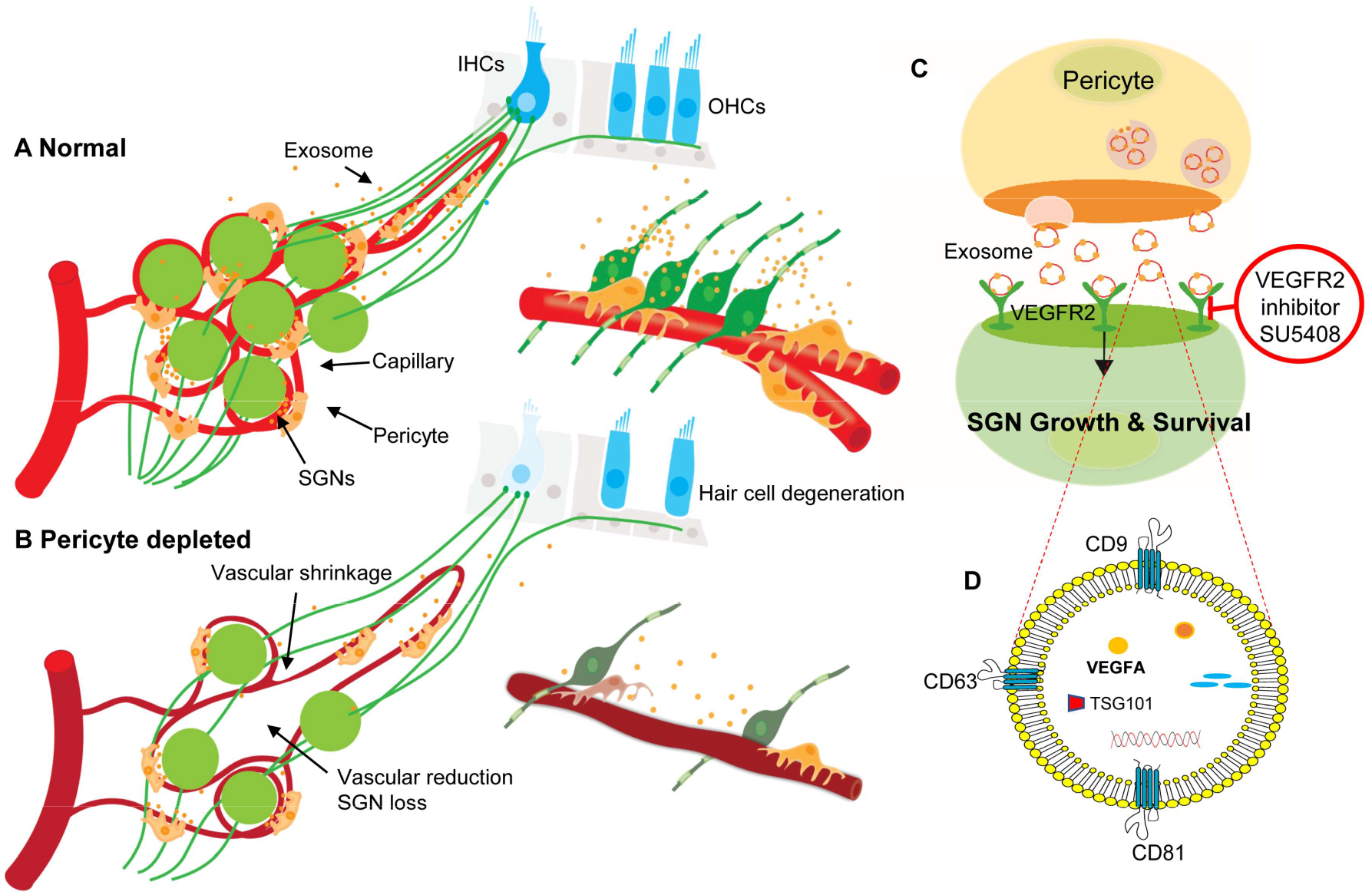
The hypothesized mechanism of pericyte-mediated SGN growth and survival. (A) Under normal conditions, pericytes participate in the maintenance of SGN health through two parallel pathways: (1) Maintenance of vascular stability and function; and (2) ‘Nourishment” of SGNs through release of exosomes. (B) Pericyte depletion causes reduction of vascular volume and dysfunction, loss of SGNs and hair cells (16), and, in consequence, hearing loss. (C) VEGF-A-carrying exosomes interact with VEGFR2 on the SGNs to stimulate growth and pro-survival response, which can be arrested by a specific VEGFR2 inhibitor, SU5408. (D) Schematic model showing the molecular structure of exosomes, including common exosome markers such as CD81, CD63, CD9 and Tsg101, and other cargo such as proteins, DNAs, RNAs, lipids and metabolites.

## Materials and Methods

### Key resources table

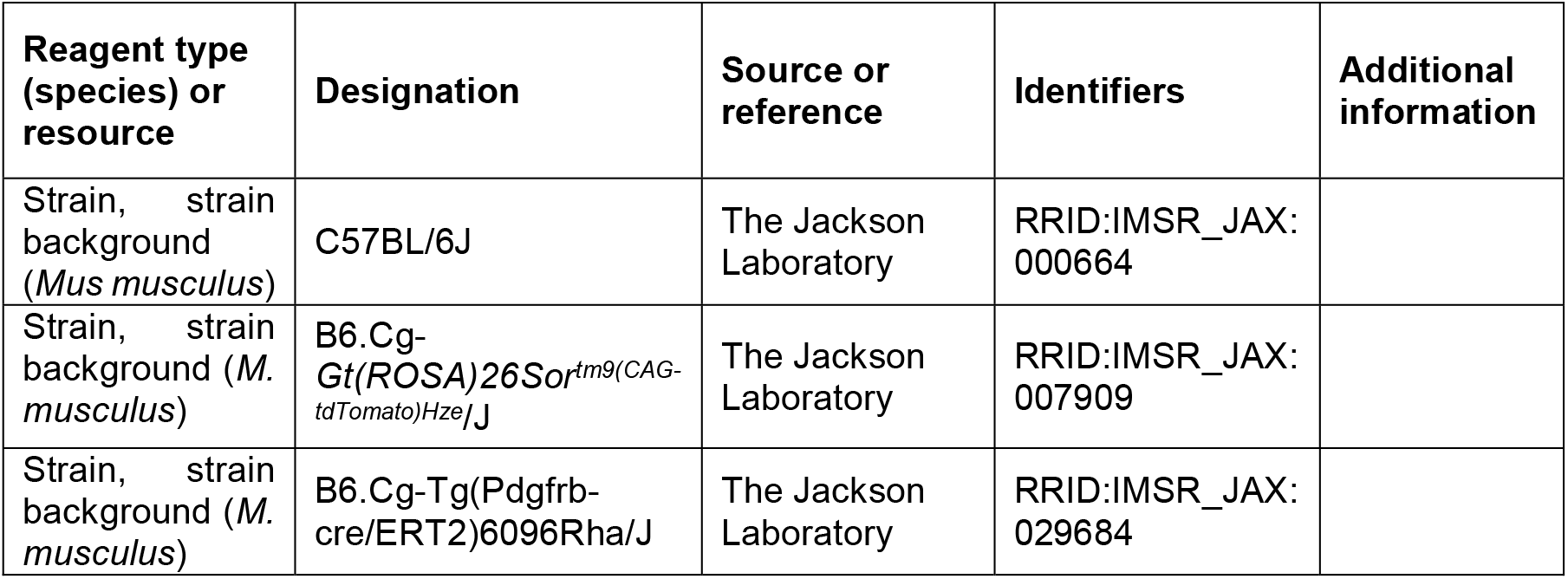
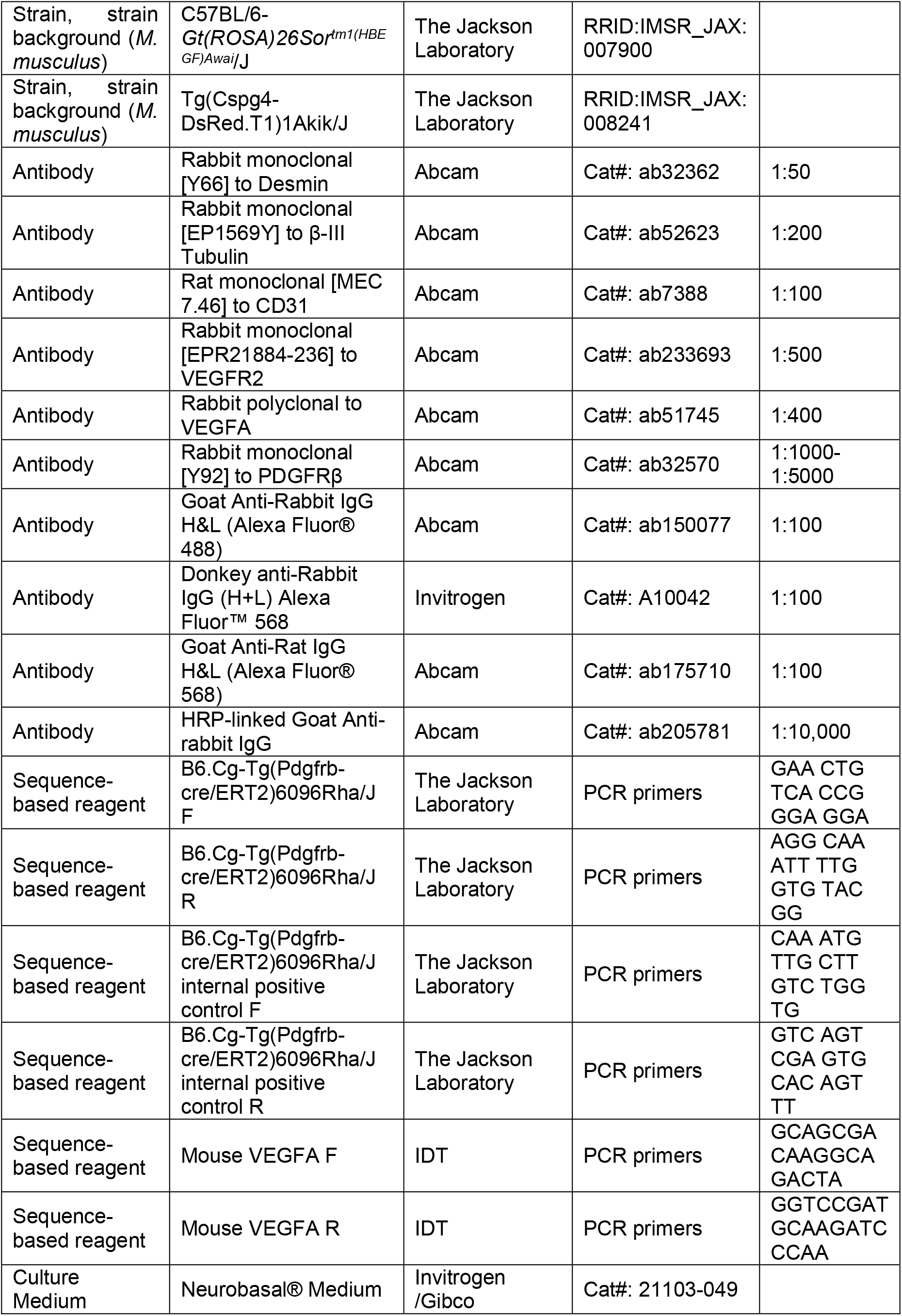
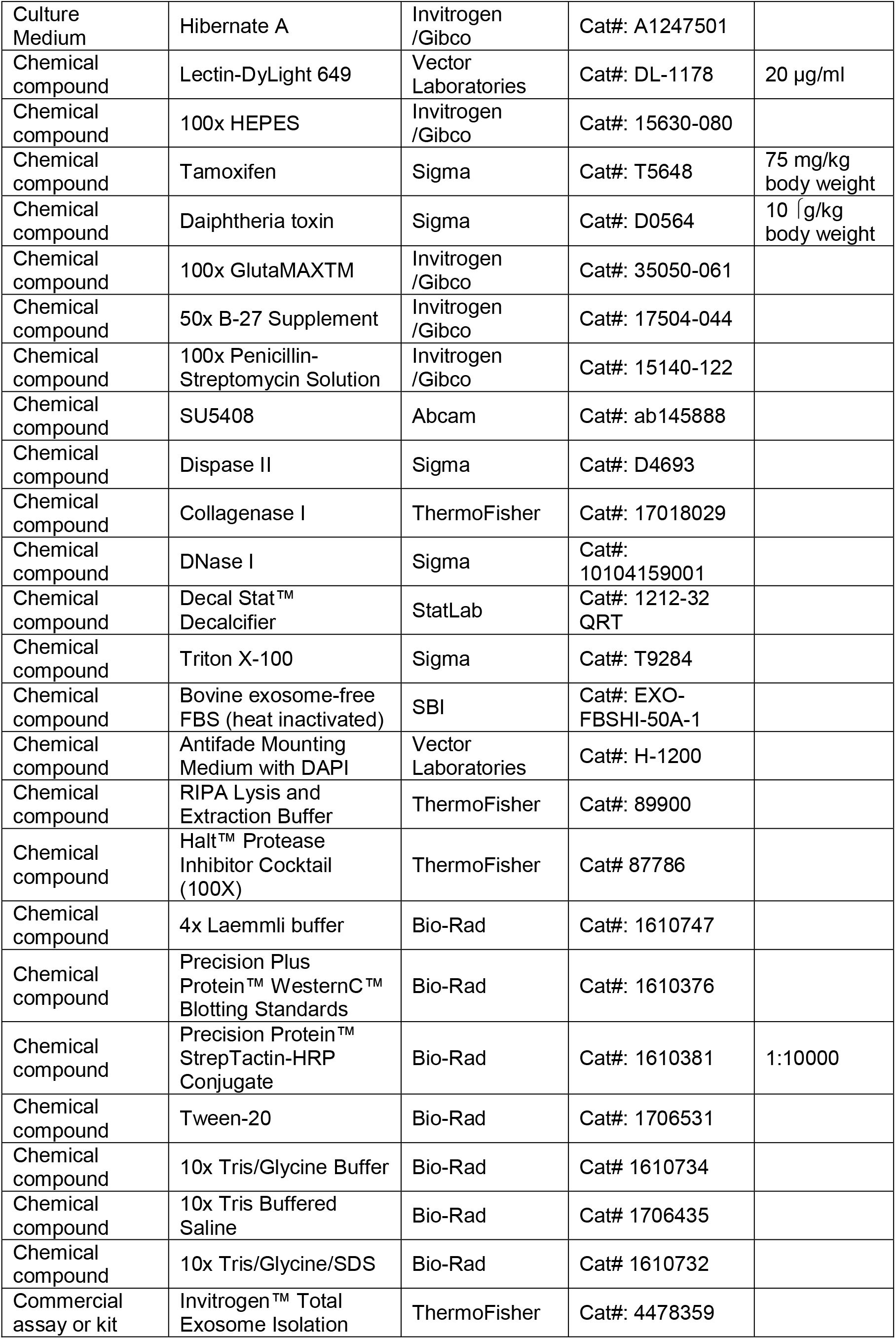
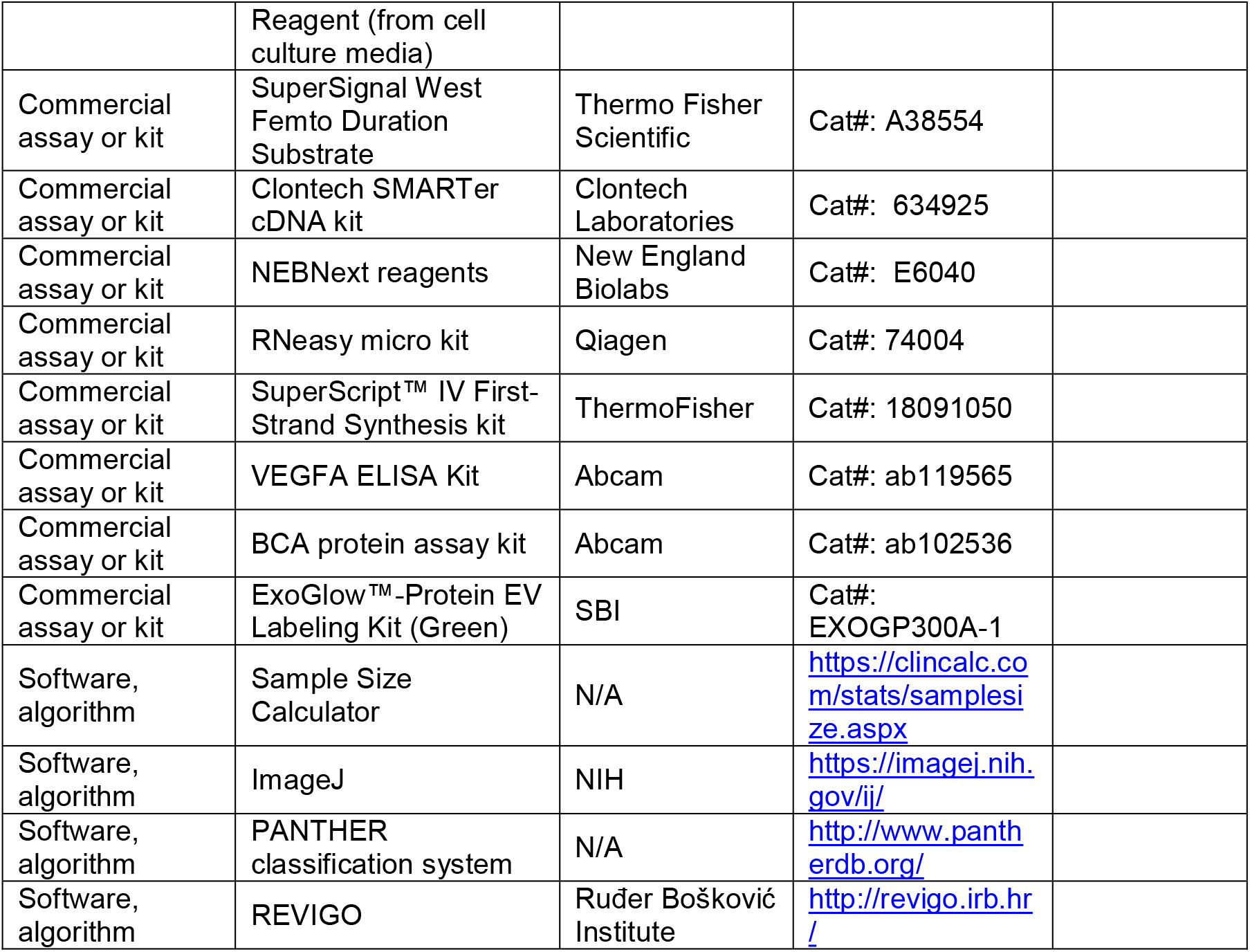

### Animals

All strains of mice used in this study were originally purchased from Jackson Laboratory, including C57BL/6J (wild type, strain # 000664), B6.Cg-*Gt(ROSA)26Sor*^*tm9(CAG-tdTomato)Hze*^/J (*ROSA26*^*tdTomato*^, strain # 007909), B6.Cg-Tg(Pdgfrb-cre/ERT2)6096Rha/J (*Pdgfrb*^*CreERT2*^, strain # 029684), C57BL/6-*Gt(ROSA)26Sor*^*tm1(HBEGF)Awai*^/J (*ROSA26*^*iDTR*^, strain # 007900), and Tg(Cspg4-DsRed.T1)1Akik/J (NG2DsRedBAC, Strain # 008241) mice. *Pdgfrb*^*CreERT2*^; *ROSA26*^*tdTomato*^ fluorescence reporter mice were created by breeding *Pdgfrb*^*CreERT2*^ mice with *ROSA26*^*tdTomato*^ mice. To verify location of the Pdgfrb/Cre, tdTomato fluorescence reporter mice were created by breeding B6.Cg-Gt (ROSA) 26Sor^tm9(CAG-tdTomato)Hze^ with B6.Cg-Tg (Pdgfrb-cre/ERT2) 6096Rha. CreERT2-mediated recombination was initiated by intraperitoneal injection of TAM at 75 mg/body weight every 24 hrs for 3 consecutive days. The inducible pericyte depletion mouse model (*Pdgfrb*^*CreERT+/-*^; *R26*^*iDTR+/-*^) by crossing *Pdgfrb*^*CreERT2+/-*^ transgenic mice with *ROSA26*^*iDTR+/+*^ mice. To deplete pericytes, this *Pdgfrb*^*CreERT+/-*^; *R26*^*iDTR+/-*^ mice were given DT intraperitoneally once every 24 hrs at a dose of 10 ng/g on four consecutive days after administering the TAM (as illustrated in **Fig. 2A and B**). The *Pdgfrb*^*CreERT2-/-*^; *R26*^*iDTR+/-*^ mice from same litters of *Pdgfrb*^*CreERT2+/-*^; *R26*^*iDTR+/-*^ mice were treated with TAM and DT as control group. All transgenic mice were maintained in the lab and were validated and genotyped for the study. Both male and female mice were used, and all mice used were adults aged between 4 ∼ 8 weeks or postnatal P1 ∼ P3. All animal experiments reported were approved by the Oregon Health & Science University Institutional Animal Care and Use Committee (IACUC IP00000968).

### Immunofluorescence of cochlea whole mount

Cochlea whole mount was dissected and stained with fluorescently tagged antibodies as described by S. C. Montgomery *et al*. (Montgomery & Cox, 2016). Briefly, the cochleae were harvested and fixed in 4% PFA overnight. After decalcification in Decal Stat™ Decalcifier overnight at 4°C, each cochlea was carefully dissected into three whole mount turns and incubated in blocking/permeabilization solution containing 0.25% Triton X-100, 10% goat serum (GS) in 1X PBS for 1 hour at room temperature (RT), then transferred into primary antibody solution (diluted in blocking/permeabilization solution) and incubated overnight at 4°C. After three times washing with 1X PBS, samples were incubated with the fluorescence-conjugated secondary antibody in the blocking/permeabilization solution for 1 hour at RT, and washed again with 1X PBS for three times, then mounted in Antifade Mounting Medium with DAPI on slides.

### Assessment of pericyte coverage and vascular density in the spiral ganglion region

Mice were anesthetized and administered the fluorescent dye Lectin-DyLight 649 diluted in 0.1 M PBS buffer to a concentration of 20 μg/ml (vol. 100 µl) via intravenous retro-orbital sinus for 10 minutes before they were sacrificed. The cochleae whole mounts were stained with desmin 1:50 and visualized under FV1000 Olympus confocal microscope with a 10x objective. The pericyte coverage and blood vessel area were calculated using Fiji (ImageJ, NIH, 1.51t) software. Pericyte distribution was defined as pericyte 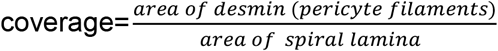. [***Note***: *Pericyte intermediate filaments were assessed to obtain pericyte coverage. We did not directly count pericyte number since all pericyte markers, including antibodies for NG2 and PDGFRβ, failed. Adult spiral limbus is very thick. For visualization of the whole mounts, including the vascular networks in the tissue, the tissues would need to be decalcified. However the decalcification process causes deterioration of membrane proteins*.]. Vascular density was analyzed as previously described (Jiang et al., 2019), defined as vascular 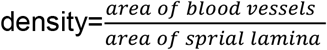.

### Demonstration of VEGFR2 expression and pericyte distribution in the spiral ganglion region

VEGFR2 expression in the region of the spiral ganglion was demonstrated in WT mice, and pericyte distribution in NG2DsRedBAC mice. Mice were anesthetized, placed on a water heating pad, and the fluorescent dye Lectin-DyLight 649 diluted in 0.1 M PBS buffer to a concentration of 20 μg/ml (vol. 100 µl) was administered via intravenous retro-orbital sinus to the animals 10 minutes before they were sacrificed. The isolated cochleae were fixed in 4% PFA overnight. The next day, tissue samples from the apical turn were isolated and the SGNs exposed after gently breaking the covering bone and removing the chips surrounding the ganglion. The tissue samples were stained with anti-VEGFR2 antibody (1:500) and/or anti-β-III tubulin (1:200) antibody, and imaged on an FV1000 Olympus laser-scanning confocal microscope. This “gentle-bone-breaking” (apex) preparation enables us to preserve the membrane protein affinity for antibodies and capture the fluorescence signal from the DsRed-tagged NG2+ pericytes.

### Auditory brainstem response (ABR) test

ABR audiometry to pure tones was used to evaluate hearing function before and after DT treatment as previously described (Zhang et al., 2021). For measurement of hearing threshold, latency, and amplitude by ABR, 10 ears from five animals in both the control and pericyte depleted group were measured by an investigator blind to treatment status. ABR response to a 1-ms rise-time tone burst (30 rep rate/sec) at 8, 16, 24, and 32 kHz was recorded. Stimulus sound pressure levels (SPL) were incremented in 5-dB steps from below threshold to 100-dB SPL. The latency at 20-100 dB and peak-to-peak (P1-N1) values for amplitudes at all tested sound pressure levels of ABR Wave I were calculated using ABR Peak Analysis 0.8 RC 1 software (Harvard-MIT Program in Speech and Hearing Bioscience and Technology).

### Immunofluorescence of cochlea frozen section

Cochleae were harvested and fixed in 4% PFA overnight. After decalcification in Decal Stat™ Decalcifier overnight at 4°C, the cochleae were dehydrated in 15% and 30% sucrose, frozen and embedded in optimal cutting temperature compound (OCT). Sections of the cochleae at 12 μm thickness were cut in the mid-modiolar plane. The specimens were permeabilized in 0.5% Triton X-100 for 30 minutes, blocked with 10% GS and 1% BSA diluted in 1X PBS for 1 hour at RT, then incubated with the primary antibodies diluted in 10% GS and 1% BSA in 1X PBS overnight at 4 °C. After three washes witih 1X PBS, the specimens were subsequently stained with fluorescence-conjugated secondary antibodies diluted in 10% GS and 1% BSA in 1X PBS for 1 hour at RT, and washed again with 1X PBS for three times, then mounted in Antifade Mounting Medium with DAPI. All samples were visualized under an FV1000 Olympus confocal microscope (Olympus FV1000, Japan).

### Assessment of SGN density and intensity

Cochleae cross-sections were stained with β-III Tubulin (1:200). Controls were prepared by replacing primary antibodies with 1% BSA-PBS. Multiple high-resolution images of each cochlea were visualized and acquired under FV1000 Olympus confocal microscope with a 40x objective. For analysis of SGNs, β-III tubulin labeled cells were visualized and counted over the same measured area of the Rosenthal’s canal. All analyses were performed with an image analysis program (Fiji). The average percentage of SGN density 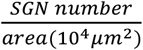 and β-III Tubulin intensity of each SGN at three regions (apex, middle and base), were calculated for each sample.

### SGN explant culture and assessment

SGN explants were isolated from P1-P3 mouse cochleae of both sexes. To maintain tissue consistency, spiral ganglion tissue from the cochlear middle turn was excised from the entire length of the cochlea and divided into three 90° fan-shaped explants, as illustrated in **Fig. 13A and B**, then cultured on a thin gel of Corning® Matrigel® Growth Factor Reduced Basement Membrane Matrix in a SGN basic culture medium consisting of Neurobasal® Medium, 1% HEPES, 1% GlutaMAXTM, 2% B-27 Supplement, and 1% Penicillin-Streptomycin Solution. [***Note:*** *serum-free medium was used to limit the effect of multi growth factors contained in the serum*]. On the final day of culture, the SGN explants and adult SGNs were fixed with 2% PFA for 30 min on ice. The specimens were permeabilized in 0.5% Triton X-100 for 30 minutes, blocked with 10% GS and 1% BSA diluted in 1X PBS for 1 hour at RT, then incubated with the antibodies for β-III tubulin (1:200) and antibody for CD31 (1:100) diluted in 10% GS and 1% BSA in 1X PBS overnight at 4 °C. After three washes witih 1X PBS, the specimens were subsequently stained with fluorescence-conjugated secondary antibodies diluted in 10% GS and 1% BSA in 1X PBS for 1 hour at RT, followed by Hoechst 33342 staining for 15 min at RT. All samples were visualized under an FV1000 Olympus confocal microscope (Olympus FV1000, Japan), as shown in **Fig. 13C and D**. Using Fiji, the total number of neurites and blood vessel sprouts (new branch formations outside the tissue explants) were counted, then divided by total area of the explant for quantification, as well as the average length of neurites assessed, 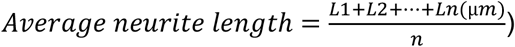 (**Fig. 13E and F**).

**Fig. 13.**
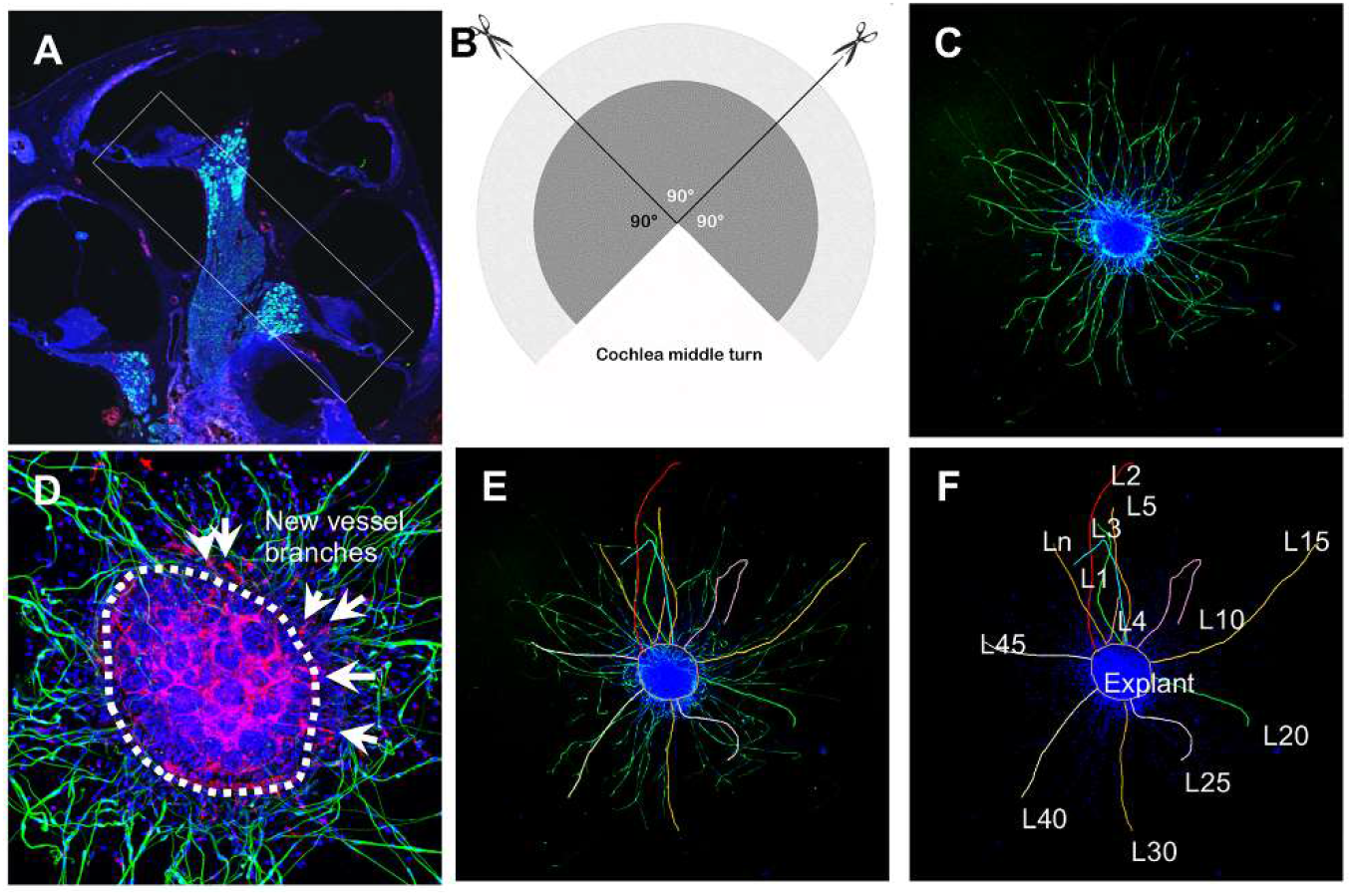
Schematic of dissection and culture of SGN explants, and method of quantitative analysis of neural dendritic growth in a P2 neonatal mouse cochlea over the course of 5 days in culture. (A and B) The cochlear middle turns were dissected out, stria vascularis and organ of Corti discarded, and the remaining SGN cut into three 90° fan-shaped pieces. These were attached to a coated 6-well glass bottom plate and cultured. (C) Representative confocal projection image of an SGN explant labeled with an antibody for β-III tubulin (green). (D-F) Illustration showing the method used for quantification of SGN neurite number and length. N: the number of neurites; L: the length of the neurites. (F) High magnification of the SGN explant showing new vessel growth labeled with CD31 (red). Arrows indicate new vessel branches.

### Adult SGNs Culture and assessment

Adult spiral ganglion neuron cells were isolated and cultured using modified protocol described by Vieira M *et al*. (Vieira et al., 2007). Briefly, cochleae from 4 weeks old mice of both sexes were collected, after the organ of Corti was peeled away, the modiolus was split longitudinally to expose the spiral ganglion and digested in 1 ml Hibernate A containing 2 mg/ml dispase II, 1 mg/ml collagenase I and 1 mg/ml DNase I for 30 min at 33 °C, gently agitated every 5 min. The tissue was dissociated by aspirating and expelling 30 times through a 1 ml polypropylene pipet tip. Tissue clumps were allowed to sediment for 1-2 min, the supernatants were collected and centrifuged at at 80 xg for 1 min to remove bony debris, the supernatant was slowly transfered to a new 15 ml centrifuge tube containing 3ml of 15% BSA+10% FBS in Hibernate A as a cushion to remove the fiber debris by centrifuging at 200 xg for 10 min. During centrifugation, the cells will pass the cushion and go down to the bottom while the fiber debris will be accumulated in the cushion layer. The pelleted cells were finally resuspended at 3 × 10^4^ neurons/ml in SGN basic culture medium (see **SGN explant culture**), ∼3000 SGN cells in 100 μl culture medium were added to each well coated with matrigel matrix diluted in neurobasal medium at a ratio of 1: 9. On the final day of culture, the adult SGNs were fixed with 2% PFA for 30 min on ice. The specimens were permeabilized in 0.5% Triton X-100 for 30 minutes, blocked with 10% GS and 1% BSA diluted in 1X PBS for 1 hour at RT, then incubated with the antibodies for β-III tubulin (1:500) diluted in 10% GS and 1% BSA in 1X PBS overnight at 4 °C. After three washes witih 1X PBS, the specimens were subsequently stained with fluorescence-conjugated secondary antibodies diluted in 10% GS and 1% BSA in 1X PBS for 1 hour at RT, followed by Hoechst 33342 staining for 15 min at RT. All samples were visualized under an FV1000 Olympus confocal microscope (Olympus FV1000, Japan). The number of SGNs per well, neurite number, and longest neurite length of each neuron were quantified using Fiji for statistical analysis.

### Pericyte co-culture with SGN explant and adult SGNs

The primary cochlea pericyte cell line was generated from C57BL/6J mouse by a well-established ‘mini-chip’ protocol previously described and published (Neng et al., 2013), and passage 3-6 were used for all the experiments. The explants or SGNs were first incubated in 100 μl basic SGN culture medium in glass bottom 6-well plates (P06G-1.5-10-F, MatTek’s) overnight at 37°C, 5% CO_2_, allowing the tissue/cell well attached to the device. The next day, the culture medium was added up to 3 ml, and pericytes at 3.0 × 10^5^ cells/well were seeded in the insert (Cat#: 353091, Corning) for 5 continuous days. The explant was first incubated in 100 μl SGN medium in glass bottom 6-well plates overnight at 37°C, 5% CO_2_. The next day, the culture medium was added up to 3 ml, and pericytes (passage 3-6) at 3.0 × 10^5^ cells/well were seeded in the insert for 5 continuous days.

### RNA-seq analysis

Pericyte specimens were submitted to Otogenetics Corporation (Norcross, GA USA) for bulk RNA-Seq assays. Briefly, total RNA was extracted from cell pellets using the E.Z.N.A. Total RNA Kit II and the integrity and purity of total RNA were assessed using Agilent Bioanalyzer and OD260/280. 1-2 μg of cDNA was generated using Clontech SMARTer cDNA kit from 100ng of total RNA, and adaptors were removed by digestion with RsaI. The resulting cDNA was fragmented using Covaris (Covaris, Inc., Woburn, MA, USA) or Bioruptor, profiled using an Agilent Bioanalyzer, and subjected to Illumina library preparation using NEBNext reagents (Cat#: E6040, New England Biolabs). The quality and size distribution of the Illumina libraries were determined using an Agilent Bioanalyzer2100. The libraries were then submitted for Illumina HiSeq2000 sequencing, used per standard operation. Paired-end 90 or 100 nucleotide reads were generated and checked for data quality using FASTQC (Babraham Institute, Cambridge, UK), and subjected to data analysis using the platform provided by DNAnexus (DNAnexus, Inc, Mountain View, CA, USA) or the platform provided by Center for Biotechnology and Computational Biology (University of Maryland, College Park, MD USA) as previously described (Trapnell et al., 2012).

### NTA, TEM, and proteomic analysis of pericyte-derived exosomes

NTA, TEM, and proteomic analyses of pericyte-derived exosomes were performed by Alpha Nano Tech. Pericytes were initially cultured in normal media in 100 mm collagen I coated petri dishes until ∼90% confluence, then rinsed with PBS, and transferred to conditioned medium containing 2% Bovine exosome-free FBS (heat inactivated) (Cat#: EXO-FBSHI-50A-1, SBI) for 24 hours. Exosomes were purified from the collected media by ultrafiltration followed by size exclusion separation, as shown in **Fig. 5A**. Briefly, the conditioned media were loaded into the pre-rinsed ultrafiltration devices (Vivaspin 20) containing a 100 kDa molecular weight cutoff (MWCO) PES membrane, and centrifuged at 3000 x g for several intervals of 30 minutes until the final volume reached 500 μl. The exosome fractions were collected with the Izon 35 nm qEV original column, and further concentrated using Amicon Ultra 2 100 kDa MWCO centrifugal filter devices.

#### NTA

The NTA analysis of exosomes labeled with Exoglow was performed with a Zetaview Quatt (Particle Metrix, Ammersee, Germany) instrument. To label exosomes with Exoglow, 2 μl of exoglow dye was added to 12 μl of reaction buffer and mixed with 14 μl of sample, and incubated for 15 min at room temperature. Dilutions were made by mixing PBS filtered through a 0.2 μm syringe filter with a corresponding volume of sample. Particle size distribution histograms were recorded in scatter and fluorescent modes.

#### TEM

Copper carbon formvar grids were glow discharged immediately prior to loading with the sample. Sample was processed undiluted. The grid was floated on a 10 μl sample drop for 15 minutes, washed twice with water by floating on the drop of water for 30 seconds, and negatively stained with 2% uranyl acetate by floating on a drop of the stain for 30 seconds. The grid was blot dried with Whatman paper and imaged on a Jeol 1230 electron microscope.

#### Proteomic Analysis

5 ug of exosomes were dried via vacuum centrifuging. Then the sample was reconstituted in 8M urea, reduced with DTT, alkylated with iodoacetamide and digested with trypsin overnight. Peptide samples were cleaned using Pierce™ Peptide Desalting Spin Columns (Cat#: 89852, ThermoFisher) and analyzed in duplicate by LC-MS/MS using a Thermo Easy nLC 1200-QExactive HF (Cat#: LC140, ThermoFisher). Proteins were identified and quantified with Proteome Discoverer 2.5 utilizing the Uniprot mouse database appended with a common contaminants database. Further data analysis was conducted in Perseus (Log2 transformation, and GOCC termannotation). Gene Ontology (GO) enrichment analysis on these proteins was performed by using the PANTHER classification system, redundancy in the lists of enriched GO terms was minimized using REVIGO, with the similarity set to 0.5.

### RT-PCR and ELISA

Total RNA of the pericytes was extracted with an RNeasy micro kit, cDNA synthesized with SuperScript™ IV First-Strand Synthesis kit, used according to the manufacturer’s instructions. RT-PCR products (Primers: forward, GCAGCGACAAGGCAGACTA; reverse, GGTCCGATGCAAGATCCCAA, 392-bp product) were analyzed by 1.5% agarose gel electrophoresis. VEGFA expression level in the supernatant of pericyte culture media and mouse cochleae homogenate were measured by using mouse VEGFA ELISA Kit per the manufacturer’s instructions.

### Western blot

Exosomes were purified from the conditioned medium using ultracentrifugation described by Breglio AM et Al. (Breglio et al., 2020). Briefly, conditioned medium was first centrifuged at 300 ×g for 10 minutes at 4°, then at 10,000 ×g for 30 minutes at 4°C. Finally, exosomes were pelleted by ultracentrifuged at 100,000 ×g for 70 minutes at 4°C. The exosome-depleted supernatant was further concentrated with a Pierce™ Protein Concentrator PES, (3K MWCO, Cat#: 88525, ThermoFisher) at 4000 ×g and 4°C. Both the concentrated non-exosomal supernatant and the purified exosomes were resuspended in RIPA Lysis and Extraction Buffer supplemented with cOmplete Mini EDTA-Free Protease Inhibitor Cocktail (Roche) and vortexed for 1 minute. Protein concentration was analyzed with a BCA protein assay kit. The remaining sample was denatured at 95°C for 5 minutes in 4× Laemmli buffer and subjected to SDS-PAGE using 4-15% Mini-PROTEAN® TGX™ Precast Protein Gels followed by protein transfer to a 0.45-μm pore PVDF membrane (Cat#: 88585, ThermoFisher). Membranes were blocked in 3% BSA in TBS with 0.1% Tween-20 (TBS-T). The primary Abs used were VEGFA (1:400) and PDGFRβ (1:5000). HRP-linked secondary Ab: Goat Anti-rabbit IgG (1:10,000). Protein bands were visualized by chemiluminescence using a SuperSignal West Femto Duration Substrate and Q-View™ Imager System (Quansys Bioscience, Logan, UT, USA).

### SGN explant and adult SGNs treatment with pericyte-derived exosomes

Exosomes were purified from the conditioned media using Invitrogen™ Total Exosome Isolation Reagent (from cell culture media) (Cat#: 4478359, ThermoFisher) according to the manufacturer’s instructions. The explants or SGNs were cultured in 400 μl SGN basic medium in Nunc™ Lab-Tek™ II Chambered Coverglass (Cat#: 155409, ThermoFisher) with exosomes (5 μg/ml), with or without SU5408 (100 nM for SGN explant, 50 nM for SGNs) for 5 continuous days. All media were changed every 2 days.

### Statistics

All experiments were designed with proper controls. The sample size estimation was conducted based on our pilot data by power analysis (Sample Size Calculator).To avoid variation and experimental bias, we applied a blind control procedure. All statistical analyses were performed using GraphPad Prism 9 software (GraphPad Software). Statistical difference between two groups was evaluated by unpaired, two-tailed t test. One-way ANOVA followed by Tukey’s multiple comparison test was used to compare differences across multiple groups. Two-way ANOVA followed by Dunnett’s multiple comparison test was used for ABR analysis to compare threshold or wave I latency differences at different time points, and wave I amplitude differences at different sound pressure levels within multiple groups. Differences were considered significant at p<0.05. Data are presented as the mean ± SEM.

## Acknowledgments

This research was supported by NIH/NIDCD R21 DC016157 (X. Shi), NIH/NIDCD R01 DC015781 (X. Shi), NIH/NIDCD R01-DC010844 (X. Shi).

## References

Abhinand, C. S., Raju, R., Soumya, S. J., Arya, P. S., & Sudhakaran, P. R. (2016). VEGF-A/VEGFR2 signaling network in endothelial cells relevant to angiogenesis. J Cell Commun Signal, 10(4), 347–354. https://doi.org/10.1007/s12079-016-0352-8

Angelborg, C., Axelsson, A., & Larsen, H.-C. (1984). Regional blood flow in the rabbit cochlea. Archives of Otolaryngology, 110(5), 297–300.

Armulik, A., Genove, G., & Betsholtz, C. (2011). Pericytes: developmental, physiological, and pathological perspectives, problems, and promises. Dev Cell, 21(2), 193–215. https://doi.org/10.1016/j.devcel.2011.07.001

Armulik, A., Genove, G., Mae, M., Nisancioglu, M. H., Wallgard, E., Niaudet, C., He, L., Norlin, J., Lindblom, P., Strittmatter, K., Johansson, B. R., & Betsholtz, C. (2010). Pericytes regulate the blood-brain barrier. Nature, 468(7323), 557–561. https://doi.org/10.1038/nature09522

Attwell, D., Mishra, A., Hall, C. N., O’Farrell, F. M., & Dalkara, T. (2016). What is a pericyte? J Cereb Blood Flow Metab, 36(2), 451–455. https://doi.org/10.1177/0271678X15610340

Baas, P. W., Rao, A. N., Matamoros, A. J., & Leo, L. (2016). Stability properties of neuronal microtubules. Cytoskeleton (Hoboken), 73(9), 442–460. https://doi.org/10.1002/cm.21286

Bang, C., & Thum, T. (2012). Exosomes: new players in cell-cell communication. Int J Biochem Cell Biol, 44(11), 2060–2064. https://doi.org/10.1016/j.biocel.2012.08.007

Bell, R. D., Winkler, E. A., Sagare, A. P., Singh, I., LaRue, B., Deane, R., & Zlokovic, B. V. (2010). Pericytes control key neurovascular functions and neuronal phenotype in the adult brain and during brain aging. Neuron, 68(3), 409–427. https://doi.org/10.1016/j.neuron.2010.09.043

Bellon, A., Luchino, J., Haigh, K., Rougon, G., Haigh, J., Chauvet, S., & Mann, F. (2010). VEGFR2 (KDR/Flk1) signaling mediates axon growth in response to semaphorin 3E in the developing brain. Neuron, 66(2), 205–219. https://doi.org/10.1016/j.neuron.2010.04.006

Bergers, G., & Song, S. (2005). The role of pericytes in blood-vessel formation and maintenance. Neuro Oncol, 7(4), 452–464. https://doi.org/10.1215/S1152851705000232

Birbrair, A. (2018). Pericyte Biology: Development, Homeostasis, and Disease. Adv Exp Med Biol, 1109, 1–3. https://doi.org/10.1007/978-3-030-02601-1_1

Breglio, A. M., May, L. A., Barzik, M., Welsh, N. C., Francis, S. P., Costain, T. Q., Wang, L., Anderson, D. E., Petralia, R. S., Wang, Y. X., Friedman, T. B., Wood, M. J., & Cunningham, L. L. (2020). Exosomes mediate sensory hair cell protection in the inner ear. J Clin Invest, 130(5), 2657–2672. https://doi.org/10.1172/JCI128867

Brown, L. S., Foster, C. G., Courtney, J. M., King, N. E., Howells, D. W., & Sutherland, B. A. (2019). Pericytes and Neurovascular Function in the Healthy and Diseased Brain. Front Cell Neurosci, 13, 282. https://doi.org/10.3389/fncel.2019.00282

Buch, T., Heppner, F. L., Tertilt, C., Heinen, T. J., Kremer, M., Wunderlich, F. T., Jung, S., & Waisman, A. (2005). A Cre-inducible diphtheria toxin receptor mediates cell lineage ablation after toxin administration. Nat Methods, 2(6), 419–426. https://doi.org/10.1038/nmeth762

Coate, T. M., Scott, M. K., & Gurjar, M. (2019). Current concepts in cochlear ribbon synapse formation. Synapse, 73(5), e22087. https://doi.org/10.1002/syn.22087

Colombo, M., Raposo, G., & Thery, C. (2014). Biogenesis, secretion, and intercellular interactions of exosomes and other extracellular vesicles. Annu Rev Cell Dev Biol, 30, 255–289. https://doi.org/10.1146/annurev-cellbio-101512-122326

D’Amore, P. A. (2007). Vascular endothelial cell growth factor-a: not just for endothelial cells anymore. Am J Pathol, 171(1), 14–18. https://doi.org/10.2353/ajpath.2007.070385

Dai, J., Su, Y., Zhong, S., Cong, L., Liu, B., Yang, J., Tao, Y., He, Z., Chen, C., & Jiang, Y. (2020). Exosomes: key players in cancer and potential therapeutic strategy. Signal Transduct Target Ther, 5(1), 145. https://doi.org/10.1038/s41392-020-00261-0

Dai, M., Yang, Y., Omelchenko, I., Nuttall, A. L., Kachelmeier, A., Xiu, R., & Shi, X. (2010). Bone marrow cell recruitment mediated by inducible nitric oxide synthase/stromal cell-derived factor-1alpha signaling repairs the acoustically damaged cochlear blood-labyrinth barrier. Am J Pathol, 177(6), 3089–3099. https://doi.org/10.2353/ajpath.2010.100340

Dias Moura Prazeres, P. H., Sena, I. F. G., Borges, I. D. T., de Azevedo, P. O., Andreotti, J. P., de Paiva, A. E., de Almeida, V. M., de Paula Guerra, D. A., Pinheiro Dos Santos, G. S., Mintz, A., Delbono, O., & Birbrair, A. (2017). Pericytes are heterogeneous in their origin within the same tissue. Dev Biol, 427(1), 6–11. https://doi.org/10.1016/j.ydbio.2017.05.001

Dragovic, R. A., Gardiner, C., Brooks, A. S., Tannetta, D. S., Ferguson, D. J., Hole, P., Carr, B., Redman, C. W., Harris, A. L., Dobson, P. J., Harrison, P., & Sargent, I. L. (2011). Sizing and phenotyping of cellular vesicles using Nanoparticle Tracking Analysis. Nanomedicine, 7(6), 780–788. https://doi.org/10.1016/j.nano.2011.04.003

Gaceb, A., Barbariga, M., Ozen, I., & Paul, G. (2018). The pericyte secretome: Potential impact on regeneration. Biochimie, 155, 16–25. https://doi.org/10.1016/j.biochi.2018.04.015

Gaceb, A., Ozen, I., Padel, T., Barbariga, M., & Paul, G. (2018). Pericytes secrete pro-regenerative molecules in response to platelet-derived growth factor-BB. J Cereb Blood Flow Metab, 38(1), 45–57. https://doi.org/10.1177/0271678X17719645

Gaceb, A., Özen, I., Padel, T., Barbariga, M., & Paul, G. (2018). Pericytes secrete pro-regenerative molecules in response to platelet-derived growth factor-BB. Journal of cerebral blood flow and metabolism: official journal of the International Society of Cerebral Blood Flow and Metabolism, 38(1), 45–57. https://doi.org/10.1177/0271678X17719645

Gaceb, A., & Paul, G. (2018). Pericyte Secretome. Adv Exp Med Biol, 1109, 139–163. https://doi.org/10.1007/978-3-030-02601-1_11

Greenhalgh, S. N., Iredale, J. P., & Henderson, N. C. (2013). Origins of fibrosis: pericytes take centre stage. F1000Prime Rep, 5, 37. https://doi.org/10.12703/P5-37

Greif, D. M., & Eichmann, A. (2014). Vascular biology: Brain vessels squeezed to death. Nature, 508(7494), 50–51. https://doi.org/10.1038/nature13217

Gyo, K. (2013). Experimental study of transient cochlear ischemia as a cause of sudden deafness. World J Otorhinolaryngol, 3(1), 1–15.

Hall, C. N., Reynell, C., Gesslein, B., Hamilton, N. B., Mishra, A., Sutherland, B. A., O’Farrell, F. M., Buchan, A. M., Lauritzen, M., & Attwell, D. (2014). Capillary pericytes regulate cerebral blood flow in health and disease. Nature, 508(7494), 55–60. https://doi.org/10.1038/nature13165

Hibino, H., Nin, F., Tsuzuki, C., & Kurachi, Y. (2010). How is the highly positive endocochlear potential formed? The specific architecture of the stria vascularis and the roles of the ion-transport apparatus. Pflugers Arch, 459(4), 521–533. https://doi.org/10.1007/s00424-009-0754-z

Jiang, H., Wang, X., Zhang, J., Kachelmeier, A., Lopez, I. A., & Shi, X. (2019). Microvascular networks in the area of the auditory peripheral nervous system. Hear Res, 371, 105–116. https://doi.org/10.1016/j.heares.2018.11.012

Jung, M. K., & Mun, J. Y. (2018). Sample Preparation and Imaging of Exosomes by Transmission Electron Microscopy. J Vis Exp(131). https://doi.org/10.3791/56482

Kalluri, R., & LeBleu, V. S. (2020). The biology, function, and biomedical applications of exosomes. Science, 367(6478). https://doi.org/10.1126/science.aau6977

Katsetos, C. D., Legido, A., Perentes, E., & Mork, S. J. (2003). Class III beta-tubulin isotype: a key cytoskeletal protein at the crossroads of developmental neurobiology and tumor neuropathology. J Child Neurol, 18(12), 851–866; discussion 867. https://doi.org/10.1177/088307380301801205

Kloner, R. A., King, K. S., & Harrington, M. G. (2018). No-reflow phenomenon in the heart and brain. Am J Physiol Heart Circ Physiol, 315(3), H550–H562. https://doi.org/10.1152/ajpheart.00183.2018

Leake, P. A., Akil, O., & Lang, H. (2020). Neurotrophin gene therapy to promote survival of spiral ganglion neurons after deafness. Hear Res, 107955. https://doi.org/10.1016/j.heares.2020.107955

Luck, R., Urban, S., Karakatsani, A., Harde, E., Sambandan, S., Nicholson, L., Haverkamp, S., Mann, R., Martin-Villalba, A., Schuman, E. M., Acker-Palmer, A., & Ruiz de Almodovar, C. (2019). VEGF/VEGFR2 signaling regulates hippocampal axon branching during development. Elife, 8. https://doi.org/10.7554/eLife.49818

Mei, X., Glueckert, R., Schrott-Fischer, A., Li, H., Ladak, H. M., Agrawal, S. K., & Rask-Andersen, H. (2020). Vascular Supply of the Human Spiral Ganglion: Novel Three-Dimensional Analysis Using Synchrotron Phase-Contrast Imaging and Histology. Sci Rep, 10(1), 5877. https://doi.org/10.1038/s41598-020-62653-0

Meyer, W., Godynicki, S., & Tsukise, A. (2008). Lectin histochemistry of the endothelium of blood vessels in the mammalian integument, with remarks on the endothelial glycocalyx and blood vessel system nomenclature. Ann Anat, 190(3), 264–276. https://doi.org/10.1016/j.aanat.2007.11.004

Mi, H., Muruganujan, A., Casagrande, J. T., & Thomas, P. D. (2013). Large-scale gene function analysis with the PANTHER classification system. Nat Protoc, 8(8), 1551–1566. https://doi.org/10.1038/nprot.2013.092

Mi, H., Muruganujan, A., Huang, X., Ebert, D., Mills, C., Guo, X., & Thomas, P. D. (2019). Protocol Update for large-scale genome and gene function analysis with the PANTHER classification system (v.14.0). Nat Protoc, 14(3), 703–721. https://doi.org/10.1038/s41596-019-0128-8

Montgomery, S. C., & Cox, B. C. (2016). Whole Mount Dissection and Immunofluorescence of the Adult Mouse Cochlea. J Vis Exp(107). https://doi.org/10.3791/53561

Nakashima, T., Suzuki, T., Iwagaki, T., & Hibi, T. (2001). Effects of anterior inferior cerebellar artery occlusion on cochlear blood flow–a comparison between laser-Doppler and microsphere methods. Hearing research, 162(1), 85–90.

Nayagam, B. A., Muniak, M. A., & Ryugo, D. K. (2011). The spiral ganglion: connecting the peripheral and central auditory systems. Hear Res, 278(1-2), 2–20. https://doi.org/10.1016/j.heares.2011.04.003

Neng, L., Zhang, W., Hassan, A., Zemla, M., Kachelmeier, A., Fridberger, A., Auer, M., & Shi, X. (2013). Isolation and culture of endothelial cells, pericytes and perivascular resident macrophage-like melanocytes from the young mouse ear. Nat Protoc, 8(4), 709–720. https://doi.org/10.1038/nprot.2013.033

Nikolakopoulou, A. M., Montagne, A., Kisler, K., Dai, Z., Wang, Y., Huuskonen, M. T., Sagare, A. P., Lazic, D., Sweeney, M. D., Kong, P., Wang, M., Owens, N. C., Lawson, E. J., Xie, X., Zhao, Z., & Zlokovic, B. V. (2019). Pericyte loss leads to circulatory failure and pleiotrophin depletion causing neuron loss. Nat Neurosci, 22(7), 1089–1098. https://doi.org/10.1038/s41593-019-0434-z

O’Farrell, F. M., & Attwell, D. (2014). A role for pericytes in coronary no-reflow. Nat Rev Cardiol, 11(7), 427–432. https://doi.org/10.1038/nrcardio.2014.58

Ogunshola, O. O., Antic, A., Donoghue, M. J., Fan, S. Y., Kim, H., Stewart, W. B., Madri, J. A., & Ment, L. R. (2002). Paracrine and autocrine functions of neuronal vascular endothelial growth factor (VEGF) in the central nervous system. J Biol Chem, 277(13), 11410–11415. https://doi.org/10.1074/jbc.M111085200

Okabe, K., Fukada, H., Tai-Nagara, I., Ando, T., Honda, T., Nakajima, K., Takeda, N., Fong, G. H., Ema, M., & Kubota, Y. (2020). Neuron-derived VEGF contributes to cortical and hippocampal development independently of VEGFR1/2-mediated neurotrophism. Dev Biol, 459(2), 65–71. https://doi.org/10.1016/j.ydbio.2019.11.016

Pfister, F., Feng, Y., vom Hagen, F., Hoffmann, S., Molema, G., Hillebrands, J. L., Shani, M., Deutsch, U., & Hammes, H. P. (2008). Pericyte migration: a novel mechanism of pericyte loss in experimental diabetic retinopathy. Diabetes, 57(9), 2495–2502. https://doi.org/10.2337/db08-0325

Quaegebeur, A., Segura, I., & Carmeliet, P. (2010). Pericytes: blood-brain barrier safeguards against neurodegeneration? Neuron, 68(3), 321–323. https://doi.org/10.1016/j.neuron.2010.10.024

Roskoski, R., Jr. (2017). Vascular endothelial growth factor (VEGF) and VEGF receptor inhibitors in the treatment of renal cell carcinomas. Pharmacol Res, 120, 116–132. https://doi.org/10.1016/j.phrs.2017.03.010

Sagare, A. P., Bell, R. D., Zhao, Z., Ma, Q., Winkler, E. A., Ramanathan, A., & Zlokovic, B. V. (2013). Pericyte loss influences Alzheimer-like neurodegeneration in mice. Nat Commun, 4, 2932. https://doi.org/10.1038/ncomms3932

Shaw, I., Rider, S., Mullins, J., Hughes, J., & Peault, B. (2018). Pericytes in the renal vasculature: roles in health and disease. Nat Rev Nephrol, 14(8), 521–534. https://doi.org/10.1038/s41581-018-0032-4

Shi, X. (2011). Physiopathology of the cochlear microcirculation. Hear Res, 282(1-2), 10–24. https://doi.org/10.1016/j.heares.2011.08.006

Shrestha, B. R., Chia, C., Wu, L., Kujawa, S. G., Liberman, M. C., & Goodrich, L. V. (2018). Sensory Neuron Diversity in the Inner Ear Is Shaped by Activity. Cell, 174(5), 1229–1246 e1217. https://doi.org/10.1016/j.cell.2018.07.007

Storkebaum, E., & Carmeliet, P. (2004). VEGF: a critical player in neurodegeneration. J Clin Invest, 113(1), 14–18. https://doi.org/10.1172/JCI20682

Supek, F., Bosnjak, M., Skunca, N., & Smuc, T. (2011). REVIGO summarizes and visualizes long lists of gene ontology terms. PLoS One, 6(7), e21800. https://doi.org/10.1371/journal.pone.0021800

Sweeney, M. D., Sagare, A. P., & Zlokovic, B. V. (2018). Blood-brain barrier breakdown in Alzheimer disease and other neurodegenerative disorders. Nat Rev Neurol, 14(3), 133–150. https://doi.org/10.1038/nrneurol.2017.188

Teran, M., & Nugent, M. A. (2019). Characterization of receptor binding kinetics for vascular endothelial growth factor-A using SPR. Anal Biochem, 564-565, 21–31. https://doi.org/10.1016/j.ab.2018.10.001

Trapnell, C., Roberts, A., Goff, L., Pertea, G., Kim, D., Kelley, D. R., Pimentel, H., Salzberg, S. L., Rinn, J. L., & Pachter, L. (2012). Differential gene and transcript expression analysis of RNA-seq experiments with TopHat and Cufflinks. Nat Protoc, 7(3), 562–578. https://doi.org/10.1038/nprot.2012.016

Viana, L. M., O’Malley, J. T., Burgess, B. J., Jones, D. D., Oliveira, C. A., Santos, F., Merchant, S. N., Liberman, L. D., & Liberman, M. C. (2015). Cochlear neuropathy in human presbycusis: Confocal analysis of hidden hearing loss in post-mortem tissue. Hear Res, 327, 78–88. https://doi.org/10.1016/j.heares.2015.04.014

Vieira, M., Christensen, B. L., Wheeler, B. C., Feng, A. S., & Kollmar, R. (2007). Survival and stimulation of neurite outgrowth in a serum-free culture of spiral ganglion neurons from adult mice. Hear Res, 230(1-2), 17–23. https://doi.org/10.1016/j.heares.2007.03.005

Winkler, E. A., Bell, R. D., & Zlokovic, B. V. (2011). Central nervous system pericytes in health and disease. Nat Neurosci, 14(11), 1398–1405. https://doi.org/10.1038/nn.2946

Xie, L., Wang, M., Liao, T., Tan, S., Sun, K., Li, H., Fang, Q., & Tang, A. (2018). The characterization of auditory brainstem response (ABR) waveforms: A study in tree shrews (Tupaia belangeri). J Otol, 13(3), 85–91. https://doi.org/10.1016/j.joto.2018.05.004

Ye, L., Guo, H., Wang, Y., Peng, Y., Zhang, Y., Li, S., Yang, M., & Wang, L. (2021). Exosomal circEhmt1 Released from Hypoxia-Pretreated Pericytes Regulates High Glucose-Induced Microvascular Dysfunction via the NFIA/NLRP3 Pathway. Oxid Med Cell Longev, 2021, 8833098. https://doi.org/10.1155/2021/8833098

Yemisci, M., Gursoy-Ozdemir, Y., Vural, A., Can, A., Topalkara, K., & Dalkara, T. (2009). Pericyte contraction induced by oxidative-nitrative stress impairs capillary reflow despite successful opening of an occluded cerebral artery. Nat Med, 15(9), 1031–1037. https://doi.org/10.1038/nm.2022

Yuan, X., Wu, Q., Wang, P., Jing, Y., Yao, H., Tang, Y., Li, Z., Zhang, H., & Xiu, R. (2019). Exosomes Derived From Pericytes Improve Microcirculation and Protect Blood-Spinal Cord Barrier After Spinal Cord Injury in Mice. Front Neurosci, 13, 319. https://doi.org/10.3389/fnins.2019.00319

Zehendner, C. M., Sebastiani, A., Hugonnet, A., Bischoff, F., Luhmann, H. J., & Thal, S. C. (2015). Traumatic brain injury results in rapid pericyte loss followed by reactive pericytosis in the cerebral cortex. Sci Rep, 5, 13497. https://doi.org/10.1038/srep13497

Zhang, J., Hou, Z., Wang, X., Jiang, H., Neng, L., Zhang, Y., Yu, Q., Burwood, G., Song, J., Auer, M., Fridberger, A., Hoa, M., & Shi, X. (2021). VEGFA165 gene therapy ameliorates blood-labyrinth barrier breakdown and hearing loss. JCI Insight, 6(8). https://doi.org/10.1172/jci.insight.143285

Zhang, Y., Liu, Y., Liu, H., & Tang, W. H. (2019). Exosomes: biogenesis, biologic function and clinical potential. Cell Biosci, 9, 19. https://doi.org/10.1186/s13578-019-0282-2

Zlokovic, B. V. (2011). Neurovascular pathways to neurodegeneration in Alzheimer’s disease and other disorders. Nat Rev Neurosci, 12(12), 723–738. https://doi.org/10.1038/nrn3114

